# *N*-Hydroxypipecolic Acid Triggers Systemic Acquired Resistance Against Leaf Rust (*Puccinia triticina*) in Wheat

**DOI:** 10.64898/2026.11.12.759722

**Authors:** Sherry Sun, Sean Formby, Caley Craven, Janesse Holmes, Cornelia Herrfurth, Lei Tian, Stefanie König, Ivo Feussner, Yuelin Zhang, Guus Bakkeren

**Affiliations:** Michael Smith Laboratories, Department of Botany, University of British Columbia, Vancouver, B.C., V6T 1Z4, Canada; Agriculture and Agri-Food Canada, Summerland Research and Development Centre, Summerland, B.C., V0H 1Z0, Canada; Bioinformatics Program, The University of British Columbia, Vancouver, BC, V5T4S6, Canada; University of Göttingen, Albrecht von Haller Institute for Plant Sciences and Göttingen Center for Molecular Biosciences (GZMB), Department of Plant Biochemistry, Göttingen, Germany; Biological Sciences Building, Department of Botany, University of British Columbia, Vancouver, B.C., V6T 1Z4, Canada

**Author notes:** The author(s) responsible for distribution of materials integral to the findings presented in this article in accordance with the policy described in the Author guidelines (https://academic.oup.com/plcell/pages/author-guidelines) is (are): Sherry Sun and Guus Bakkeren.

**Keywords:** Systemic acquired resistance (SAR), Phytopathogenesis, fungal pathogen, biotrophy, wheat leaf rust disease, multi-omics, phytohormones

## Abstract

Systemic acquired resistance (SAR) represents a conserved immune strategy enabling plants to mount long-lasting, whole-plant protection following localized pathogen challenge. In *Arabidopsis thaliana*, SAR is orchestrated by the mobile immune regulator *N-*hydroxypipecolic acid (NHP); however, the extent to which NHP defines systemic immune states in polyploid crops remains unclear. Here, we establish NHP as an inducer of systemic immunity in hexaploid wheat (*Triticum aestivum*) during infection by the obligate biotrophic fungus *Puccinia triticina*. Local NHP application conferred robust disease suppression in distal leaves and elicited a temporally structured oxidative response characterized by a distal, biphasic reactive oxygen species (ROS) wave linking local and systemic tissues. RNA-sequencing analysis revealed that NHP induces coordinated transcriptional reprogramming, characterized by systemic activation of pattern-triggered and effector-triggered immunity pathways, WRKY-centered transcriptional networks, NPR1-associated signaling, and broad-spectrum resistance loci including *Lr34*. Concomitantly, targeted metabolomic profiling revealed a unique systemic accumulation of salicylic acid, salicylic acid glucoside, pipecolic acid, and NHP, without the conventional jasmonate signaling antagonism observed in eudicots. Despite this conserved immune architecture, functional analyses indicated divergence within the NHP biosynthetic pathway; while a single wheat *FMO* from sub-clade A2 was systemically up-regulated by NHP, the highly conserved *TaSARD4* homeologs were constitutively expressed and functionally dispensable for rust resistance *in planta*. Notably, heterologous expression of *TaSARD4-B* successfully restored NHP-glucoside accumulation in an Arabidopsis *sard4* mutant, confirming biochemical competence and suggesting alternative metabolic redundancies in wheat. These findings define a systems-level NHP-driven immune state in wheat and provide a conceptual framework for dissecting systemic immunity in cereals, extending SAR paradigms beyond model plant systems.

**Short summary:** *N*-hydroxypipecolic acid (NHP) is a central regulator of systemic acquired resistance in Arabidopsis, yet its role in cereal crops remains unclear. We show that exogenous NHP induces a coordinated systemic immune state in hexaploid wheat, reducing susceptibility to the biotrophic fungus *Puccinia triticina*. Integrated histological, metabolomic, and transcriptomic analyses reveal conserved SAR hallmarks alongside cereal-specific regulatory features, defining an NHP-induced resistance (NIR) program in wheat and expanding our understanding of systemic immunity in cereal crops.

## Introduction

Bread wheat (*Triticum aestivum* L., *Ta*) is a critical global dietary staple (Hawkesford et al., 2013; Shewry, 2009; Venske et al., 2019), yet its production is consistently threatened by destructive fungal diseases (Savary et al., 2019; J. Singh et al., 2023). Among the globally most economically devastating is wheat leaf rust, caused by the obligate biotrophic fungus *Puccinia triticina* (*Pt*), which drains host nutrients and drastically reduces grain yield (Bakkeren et al., 2016; Bolton et al., 2008; Duplessis et al., 2014; Roelfs et al., 1992). To defend against such invasions, plants rely on a multi-tiered innate immune system. Pattern-triggered immunity (PTI) relies on cell-surface receptors to detect conserved pathogen signatures (Boller & Felix, 2009; Macho & Zipfel, 2014), while effector-triggered immunity (ETI) utilizes intracellular receptors (NLRs) to recognize pathogen-secreted virulence proteins, often culminating in localized programmed cell death (Jones et al., 2016; Jones & Dangl, 2006).

Beyond these localized defense responses, plants can also establish a long-lasting, whole- plant immune state known as systemic acquired resistance (SAR), which provides broad- spectrum protection against subsequent secondary infections in distal, uninfected tissues (Conrath, 2006; Durrant & Dong, 2004; Klessig et al., 2018; Ryals et al., 1996). SAR is widely conceptualized as a multi-phase process involving local pathogen perception (‘stage 1’), generation of mobile immune signals (‘stage 2’), their long-distance transmission and perception (‘stage 3’), and the eventual manifestation of broad-spectrum disease resistance in distal, uninfected tissues (‘stage 4’) (Cameron et al., 1994; Champigny et al., 2013; Maldonado et al., 2002). Expanding on this, the systemic propagation of SAR is considered as a highly coordinated network of parallel but interconnected signaling pathways; disparate mobile signals cooperate and fortify each other via reciprocal positive feedback loops, launching the entire plant into an ‘alarmed’ or primed state against diverse pathogens (Vlot et al., 2021).

In the eudicot model *Arabidopsis thaliana*, the establishment of SAR is orchestrated by the synergistic accumulation of salicylic acid (SA) and the lysine-derived specialized metabolite *N*-hydroxypipecolic acid (NHP) (Hartmann et al., 2018; W. Huang et al., 2020; Zeier, 2021). Upon pathogen perception, the aminotransferase AGD2-LIKE DEFENSE RESPONSE PROTEIN 1 (ALD1) converts L-lysine to a cyclic ketimine intermediate, which is subsequently reduced by SAR-DEFICIENT 4 (SARD4) to pipecolic acid (Pip); finally, the flavin-dependent monooxygenase enzyme, FLAVIN-DEPENDENT MONOOXYGENASE 1 (FMO1) hydroxylates Pip to form bioactive NHP (Figure 1a) (Y.-C. Chen et al., 2018; Ding et al., 2016; Hartmann et al., 2018; Song et al., 2004). Once synthesized, NHP functions as a critical immune amplifier and mobile signal, driving the systemic accumulation of SA, massive transcriptional reprogramming via the NPR1 regulatory hub, and the potentiation of reactive oxygen species (ROS) bursts necessary to restrict pathogen colonization (Y.-C. Chen et al., 2018; Kachroo & Kachroo, 2018; Tian & Zhang, 2019; Yildiz et al., 2021; Zeier, 2021; P. Zhang et al., 2026).

**Figure 1.**
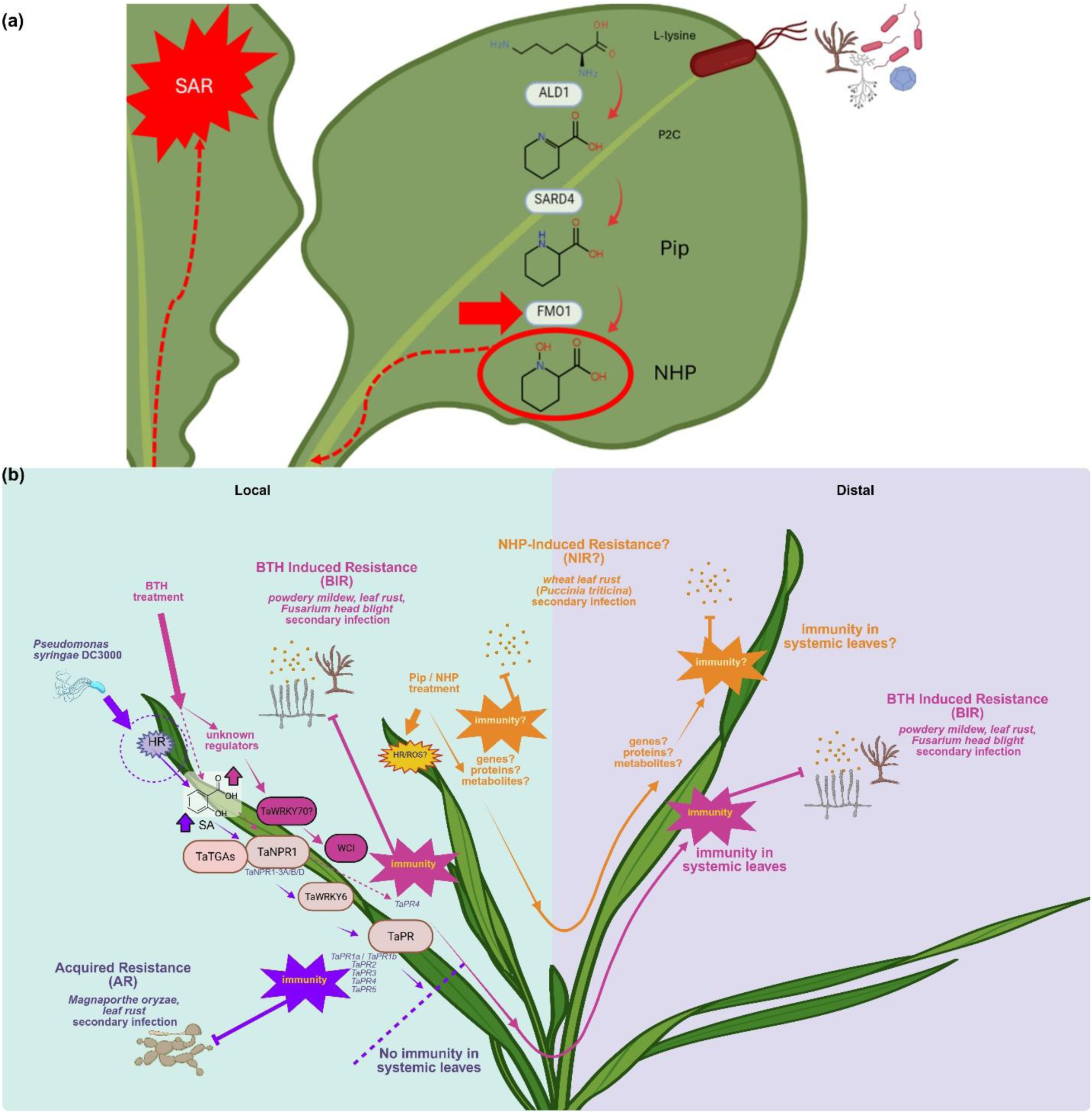
The canonical Arabidopsis NHP biosynthetic pathway and established models of induced resistance in hexaploid wheat. **(a)** NHP biosynthesis and systemic acquired resistance (SAR) in the eudicot model *Arabidopsis thaliana*. Upon pathogen perception, L-lysine is converted to a cyclic Δ^1^-piperideine-2-carboxylic acid (P2C) intermediate by the aminotransferase ALD1. P2C is subsequently reduced to pipecolic acid (Pip) by the ketimine reductase SARD4, and finally hydroxylated into the bioactive mobile signal *N*-hydroxypipecolic acid (NHP) by the flavin-dependent monooxygenase FMO1. The accumulation of NHP establishes SAR, conferring broad-spectrum protection in distal, uninfected tissues. **(b)** Current models of systemic-like immunity in wheat and the proposed NHP-Induced Resistance (NIR) pathway. Historically, induced disease resistance in cereals comprises two partially overlapping pathways. Acquired Resistance (AR, purple arrows) is triggered by localized bacterial infection (e.g., *Pseudomonas syringae* DC3000), driving local salicylic acid (SA) accumulation and the activation of *TaNPR1* homeologs and *TaTGA*s. This up-regulates *TaWRKY6* and *TaPR* genes to restrict secondary infections (such as *Magnaporthe oryzae* or *Puccinia triticina*) at the local site but does not confer systemic immunity. Benzothiadiazole (BTH)-Induced Resistance (BIR, pink arrows) is triggered by the SA-analog BTH, engaging *TaNPR1* and other unknown regulators to activate *TaWRKY70* and *Wheat Chemically Induced* (*WCI*) genes, establishing whole-plant systemic immunity against powdery mildew, leaf rust, and *Fusarium* head blight. The proposed NHP-Induced Resistance pathway (NIR, orange arrows) illustrates the core knowledge gap addressed in this study: whether local supplementation of Pip or NHP can successfully trigger early reactive oxygen species (ROS) networks, whole-plant transcriptional reprogramming, and systemic immunity against the obligate biotrophic rust fungus *Pt*. Solid arrows denote established relationships; dashed arrows represent inferred or unknown signaling mechanisms. **Alt text:** Schematic diagram illustrating the Arabidopsis NHP biosynthetic pathway alongside current models of Acquired Resistance, BTH-Induced Resistance, and proposed NHP-Induced Resistance in wheat.

While the Pip–NHP–SA metabolic axis is well-characterized in eudicots, its conservation and functional architecture in complex polyploid cereals remain poorly defined. Historically, induced disease resistance in cereals has been described through two prominent overlapping pathways, such localized acquired resistance (AR) and benzothiadiazole (BTH)-induced resistance (BIR), the latter of which relies heavily on WRKY transcription factor networks rather than the canonical NPR1–PR pathways dominant in *Arabidopsis* (Figure 1b) (Görlach et al., 1996; H. Li et al., 2020; X. Wang et al., 2018; S. Zhao et al., 2024).

Recent evidence has detected Pip and NHP accumulation following pathogen challenge in grasses like barley and *Brachypodium* (Lenk et al., 2019; Schnake et al., 2020), and exogenous NHP has been shown to modestly enhance wheat seedling resistance against the necrotrophic fungus *Fusarium graminearum* (E. T. Zhang et al., 2021). However, it remains unknown whether exogenous NHP is sufficient to trigger full systemic immunity against obligate biotrophic rust fungi in a highly susceptible wheat host. Furthermore, the genetic architecture regulating NHP biosynthesis in cereals is highly complex; while *SARD4* orthologs are conserved, the hexaploid wheat genome harbors massive lineage-specific expansions of the FMO gene family, numbering roughly 170 candidates, obscuring the identification of functional FMO1-like orthologs (S. Sun & Bakkeren, 2024a).

In this study, we investigated the efficacy and underlying molecular mechanisms of NHP-induced systemic resistance in wheat during early *Pt* pathogenesis. By integrating histological phenotyping, global transcriptomics, and metabolite profiling, we demonstrate that local NHP application is sufficient to prime a robust, temporally coordinated systemic ROS burst and orchestrate massive transcriptional reprogramming in distal, uninfected tissues. We reveal that NHP-induced resistance (NIR) in wheat NIR in wheat is associated with a synergistic Pip–NHP–SA metabolic axis, extensive WRKY transcription factor induction, and up-regulation of broad-spectrum resistance genes, including *Lr34*. Furthermore, through *in planta* silencing and heterologous complementation of wheat *SARD4* homeologs alongside *FMO* transcriptomic mapping, we highlight key evolutionary divergences in the NHP biosynthetic machinery between monocots and eudicots.

## Materials and Methods

### Plant Material and Growth Conditions

Wheat (*Triticum aestivum* cv. Thatcher) and *Nicotiana benthamiana* plants were grown in soil in environmentally controlled growth chambers under a 16-h light/8-h dark photoperiod at 20°C (wheat) or 25°C (tobacco) with 60% relative humidity. *Arabidopsis thaliana* wild-type (Columbia-0, Col-0), *fmo1-1*, and *sard4* knock-out mutants and other derived transgenic lines were grown under a 16-h light/8-h dark photoperiod at 23°C/19°C (day/night) with 70% relative humidity.

### Pathogen Inoculation and Chemical Treatments

All fungal infection assays utilized *Pt* race 1 BBBD. Four-leaf stage wheat seedlings were locally infiltrated (second true leaf) with 10 mM *N*-hydroxypipecolic acid (NHP; Cat. No. HY-N7378, MedChemExpress, Monmouth Junction, NJ, USA), or a sterile water mock control. At 24 hours post-infiltration (hpi), whole plants were systemically spray-inoculated with *Pt* urediniospores suspended in Bayol F mineral oil (Bayol F, CAS 8042-47-5; Supelco, Bellefonte, PA, USA) at (2.0 × 10^6^ spores/mL, and incubated in a dark humidity chamber for 16 hours. Disease symptoms on distal leaves (third true leaf) were assessed at 14 days after infection (DAI) using the Fiji software plugins PIDIQ (for chlorosis) (Laflamme et al., 2016) and RUST (for pustule counts and sizes) (Gallego-Sánchez et al., 2020). Relative fungal biomass was quantified using digital droplet PCR (ddPCR; BioRad) to calculate the ratio of the *PtRTP1* fungal effector gene to the wheat *TaEF1α* reference gene.

### Histological Assessments of ROS and Fungal Structures

To visualize hydrogen peroxide (H_2_O_2_) accumulation, leaf segments were stained with 1 mg/mL 3-3′diaminobenzidine (DAB, pH 3.8; Cat# 32750; Sigma-Aldrich, St. Louis, MO, USA) (Carril et al., 2020). Fungal infection structures were co-stained using 0.1% (w/v) Uvitex 2B (Cat# NC9859220; Fisher Scientific, Waltham, MA, USA), following tissue fixation and clearing in lactophenol/ethanol (1:2 v/v). Plant cell walls were counterstained with 2 μg/mL acridine orange. Imaging was conducted using an Axiocam 512 camera and ZEN Pro software (Carl Zeiss) with Brightfield and DAPI filters. Relative DAB staining intensity was quantified across time points using the Color Deconvolution 2 macro in Fiji (Landini et al., 2021).

### RNA-Sequencing and Transcriptome Analysis

Total RNA was extracted from local and distal wheat leaf tissues using the RNeasy Plant Mini Kit (Qiagen) with on-column DNase I treatment. Libraries were prepared using the IDT xGen RNA Library Kit and sequenced on the Illumina NovaSeq X-10B platform (150-bp paired- end). Following adapter trimming and quality filtering via fastp (Chen et al., 2018), reads were pseudo-aligned to the *Ta* cv. Thatcher reference transcriptome (Li et al., submitted; https://phytozome-next.jgi.doe.gov/info/991) using Kallisto (Bray et al., 2016). Differential gene expression (DGE) was performed using the DESeq2 package in R, utilizing an additive model design (Love et al., 2014). Gene Set Enrichment Analysis (GSEA) and Gene Ontology (GO) over-representation analysis were conducted using the clusterProfiler package (Wu et al., 2021; Yu et al., 2012). Weighted Gene Co-expression Network Analysis (WGCNA) was employed to identify highly connected gene modules and hub genes associated with NHP treatment (Langfelder & Horvath, 2008). Raw sequencing reads have been deposited in the NCBI Sequence Read Archive (SRA) under the BioProject accession PRJNA1285943. All associated data matrices and R scripts utilized for these transcriptomic analyses are publicly available in the GitHub repository.

### Phylogenetic Sequence Analysis

Protein sequences of Thatcher FMO (*TaTC-FMO*) and SARD4 homologs were identified utilizing a local Hidden Markov Model (HMM) (Eddy, 2011) search alongside liftoff mapping from the *Ta* cv. Chinese Spring reference (Shumate & Salzberg, 2021). Multiple sequence alignments spanning disparate plant taxa were generated using MAFFT (Katoh et al., 2019). Maximum likelihood phylogenies were constructed using IQ-TREE (Nguyen et al., 2015), applying the Bayesian Information Criterion-selected JTT+G4 substitution model with 1000 standard bootstrap replicates (Kalyaanamoorthy et al., 2017).

### Transgenic Arabidopsis Complementation

The coding sequences of wheat homeologs *TaSARD4-A*, *-B*, and *-D* were cloned into a modified pCAMBIA1305 binary vector (pBASTA-35S-3HA-E9t) to generate C-terminal 3×HA fusion constructs. Constructs were transformed into *A. tumefaciens* strain GV3101 or EHA105 and introduced into the Arabidopsis *sard4* mutant background via the floral dip method. Transgenic lines were selected using glufosinate-ammonium (BASTA) and verified via anti-HA immunoblotting.

### Virus-Induced Gene Silencing (VIGS)

A 401-bp conserved fragment targeting all three *TaSARD4* homeologs was cloned into the Barley Stripe Mosaic Virus (BSMV) RNAγ vector (pCa-γbLIC). Following *A. tumefaciens* C58-mediated propagation in *N. benthamiana*, viral sap was rub-inoculated onto two-leaf stage wheat seedlings. Successful viral infection was monitored using a *TaPDS* positive silencing control. At 15 days post-inoculation, plants were challenged with *Pt* urediniospores, and target gene knockdown was confirmed via RT-qPCR using *TaCDC48* as an internal reference.

### Targeted Metabolomics

For Arabidopsis complementation assays, local leaves of transgenic lines were syringe-infiltrated with 100 μM NLP20 peptide to induce PTI-associated metabolite accumulation. Tissue was harvested at 24 hpi, extracted via a modified methyl-tert-butyl ether (MTBE) protocol with internal standards, reversed-phase separated using an ACQUITY UPLC® system (Waters Corp., Milford, MA, USA) and analyzed by nanoelectrospray (nanoESI) (TriVersa Nanomate^®^; Advion BioSciences, Ithaca, NY, USA) coupled with an AB Sciex 4000 QTRAP^®^ tandem mass spectrometer (AB Sciex, Framingham, MA, USA) employed in scheduled multiple reaction monitoring mode for quantification of NHP-*O*Glc (Mohnike et al., 2021). For wheat assays, local and distal leaves from NHP- or mock-treated plants were harvested at 24 and 48 hpi. Phytohormones (Pip, NHP, SA, SAG, JA) were extracted using a modified QuEChERS protocol (Duszyn et al., 2022) followed by solid-phase extraction (SPE) cleanup. Quantification was performed using an ACQUITY UHPLC system coupled to a QTOF mass spectrometer (Waters), employing both positive and negative electrospray ionization modes. In wheat leaves, metabolite concentrations were determined using calibration curves generated from authentic external standards and normalized to sample fresh weight.

### Statistical Analyses

Statistical analyses were performed in R (v4.4.1). Phenotypic and histological comparisons between treatments were analyzed using independent Student’s t-tests. Phytohormone and metabolite data were subjected to a two-way analysis of variance (ANOVA, Type II test) followed by Bonferroni or Tukey HSD post-hoc pairwise comparisons.

## Results

### Local NHP treatment enhances systemic resistance to wheat leaf rust

To establish the biological relevance of *N*-hydroxypipecolic acid (NHP)–mediated signaling in wheat, we first asked whether local application of NHP was sufficient to induce systemic resistance against the leaf rust pathogen *Pt*. Four-leaf stage seedlings of the susceptible wheat cultivar Thatcher were locally infiltrated with either a mock water treatment or 10 mM NHP. 24 hours after this local pre-treatment, the plants were systemically spray-inoculated with *Pt* race 1 BBBD urediniospores, and disease symptoms and fungal colonization were assessed in the distal, untreated leaves at 14 days after infection (DAI).

Local NHP treatment 24 hours prior to whole-plant *Pt* inoculation triggered a pronounced systemic protective effect in distal, untreated leaves relative to mock controls, characterized by visibly reduced pustule formation and disease-associated chlorosis (Figure 2c). Plants receiving local 10 mM NHP exhibited a significant 3.5-fold reduction in infection frequency (IF; ∼14.4 vs. ∼50.5 pustules/cm^2^, *p* < 0.05) (Supplementary Figure 1a) and a 4-fold decrease in total disease coverage (∼3.8% vs. ∼15.3%, *p* < 0.01) (Supplementary Figure 1b). Consistent with these phenotypic observations, digital droplet PCR (ddPCR) quantification of fungal biomass revealed a significant, approximately 2-fold reduction (*p* < 0.01) in pathogen accumulation in systemic tissue following NHP treatment. This reduction in fungal biomass was sustained from 7 to 14 DAI, indicating durable systemic protection (Figure 2a,b).

**Figure 2.**
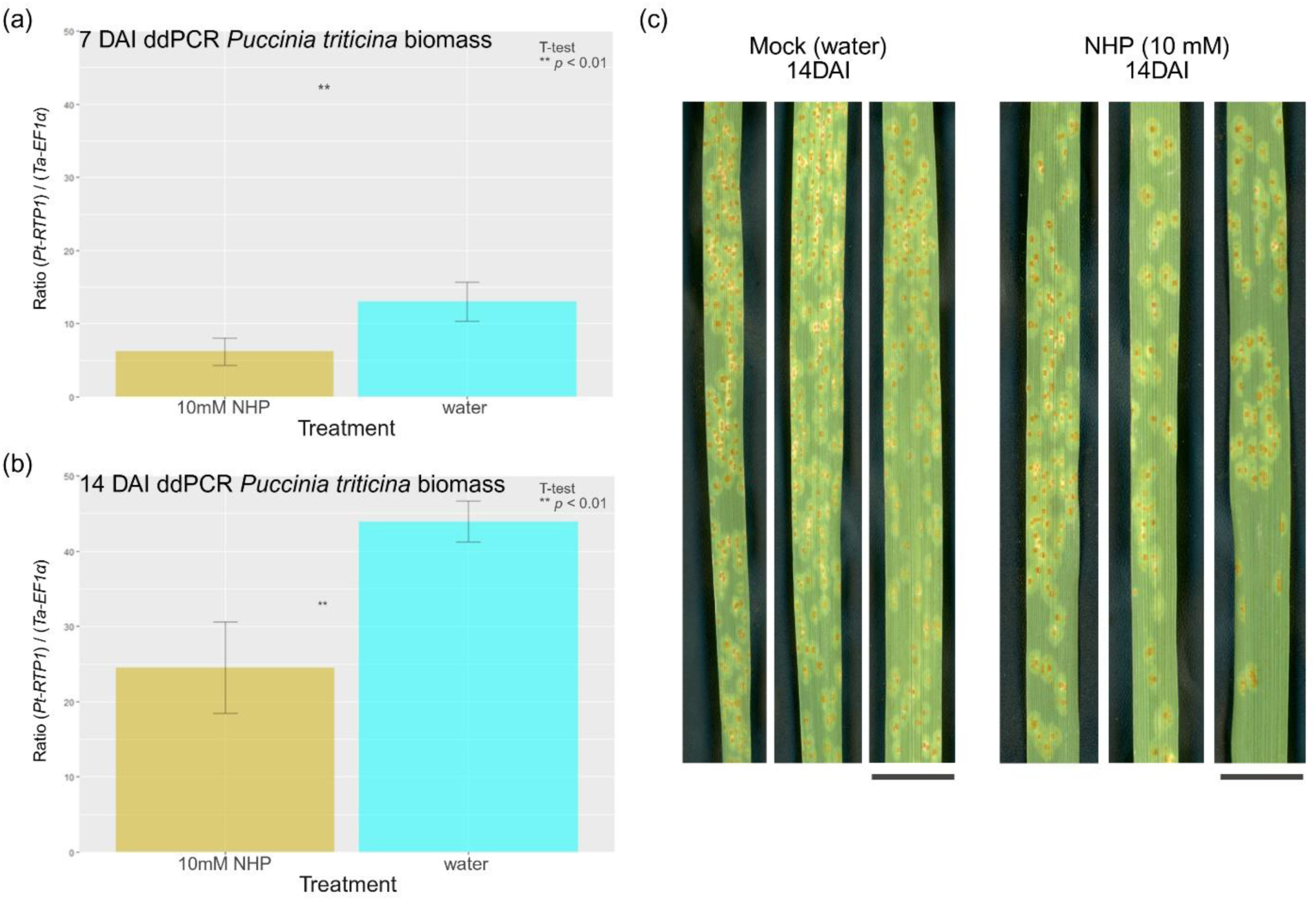
Local *N*-hydroxypipecolic acid (NHP) pre-treatment restricts *Puccinia triticina* fungal biomass and systemically reduces disease symptoms in susceptible wheat. Four-leaf stage susceptible wheat plants (cv. Thatcher) were locally infiltrated in the third leaf with a mock treatment (water) or 10 mM NHP. At 24 hours post-treatment, whole plants were systemically infected with *Puccinia triticina* (*Pt*, race 1 BBBD) urediniospores. (a–b) Relative *Pt* fungal biomass in distal (untreated) leaves of NHP-treated versus mock-treated plants at (a) 7 days after infection (DAI) and (b) 14 DAI, quantified via digital droplet PCR (ddPCR) as the ratio of the fungal single-copy effector gene *PtRTP1* to the wheat single-copy reference gene *TaEF1α*. Bar plots represent mean values ± the standard error of the mean (SEM) from three independent experiments (*n* = 3 biological replicates), with each replicate comprising pooled samples from eight or nine individual plants. Asterisks indicate statistically significant differences determined by independent two-tailed Student’s t-tests: ** *p* < 0.01. (c) Representative line-up of distal leaves at 14 DAI comparing mock- and NHP-treated plants. **Alt text:** Bar plots and leaf photos showing local 10 mM NHP pre-treatment significantly reduces distal *Puccinia triticina* fungal biomass and rust disease symptoms at 7 and 14 days post-infection.

Together, these results demonstrate that local application of exogenous NHP is sufficient to induce robust and sustained systemic resistance and delimit fungal colonization in an otherwise susceptible wheat-*Pt* interaction. This establishes a functional foundation for dissecting the downstream regulatory mechanisms governing NHP-mediated SAR-like immunity in a complex cereal genome.

### NHP application augments a temporally resolved systemic ROS burst at *Pt* infection sites

Systemic resistance is preceded by a coordinated reactive oxygen species (ROS) wave, consistent with SAR-like signaling. To confirm the systemic defense initiation observed phenotypically, we assessed the production of ROS, specifically hydrogen peroxide H_2_O_2_, as an early cellular marker of defense priming. Four-leaf stage wheat seedlings were locally pre- treated with either 10 mM NHP or water, followed by systemic *Pt* inoculation at 24 hours post- treatment. Leaf segments were co-stained with 3,3′-diaminobenzidine (DAB) to quantify relative H_2_O_2_ accumulation and Uvitex 2B to visualize fungal penetration structures at 24 and 48 hours post-infection (hpi) (Figure 3a,b).

**Figure 3.**
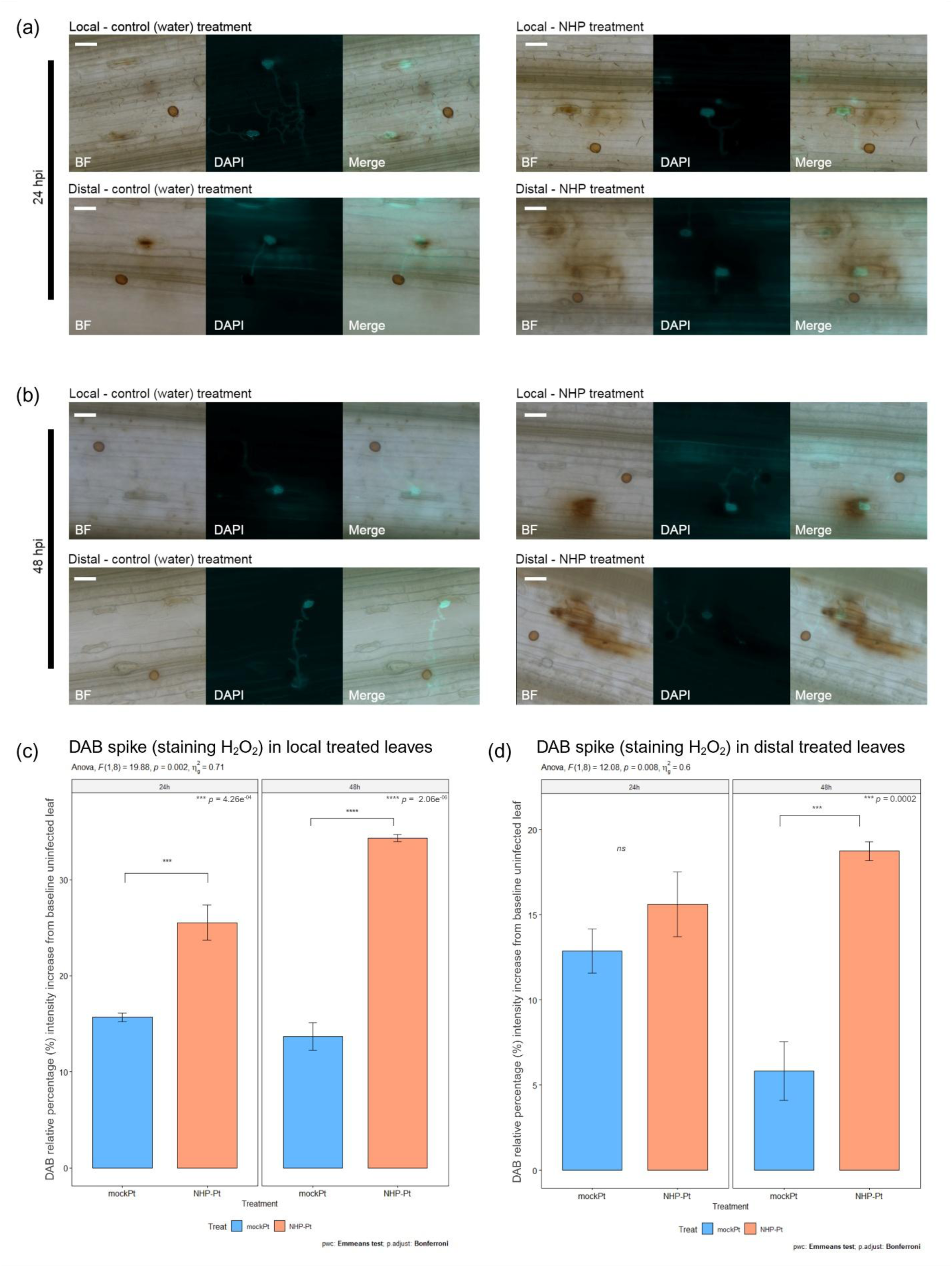
NHP induces a sustained local H₂O₂ burst and a delayed systemic ROS wave during early *Pt* infection. Four-leaf stage Thatcher wheat plants were locally infiltrated in the third leaf (second true leaf) with either a mock treatment (water) or 10 mM NHP. At 24 hours post-treatment, plants were systemically inoculated with *Pt* race 1 BBBD urediniospores. Local and distal leaves were co-stained with DAB (to visualize H₂O₂ accumulation) and Uvitex 2B (to visualize fungal penetration structures). **(a, b)** Representative micrographs of local and distal leaves from mock-treated and NHP-treated plants at 24 hpi **(a)** and 48 hpi **(b)**. The Brightfield (BF) channel captures the leaf surface, stomata, and brown DAB precipitate; the DAPI channel visualizes Uvitex 2B-stained *Pt* infection structures (germination tubes, penetration pegs); and the Merge panel shows superimposed channels. Scale bars = 50 μm. **(c, d)** Quantification of relative DAB staining intensity in local treated leaves **(c)** and distal untreated leaves **(d)** at 24 and 48 hpi. Data are expressed as the relative percentage increase in DAB intensity compared to a blank, uninfected control leaf. Bar plots represent mean values (± SEM) from three independent experiments (*n* = 3 biological replicates), with each replicate pooling leaves from eight individual plants per treatment. Statistical significance between mock (blue) and NHP (orange) treatments at each time point was determined via two-way ANOVA followed by Bonferroni- adjusted post hoc comparisons (*** *p* < 0.001, **** *p* < 0.0001, ns = not significant). **Alt text:** Micrographs and bar plots illustrating local DAB-stained H_2_O_2_ accumulation at 24-48 hpi and systemic ROS elevation at 48 hpi following NHP treatment and *Pt* infection.

Local NHP administration triggered a robust and sustained ROS burst at the sites of infection. In local leaves, DAB staining was highly concentrated around stomata penetrated by *Pt* infection pegs (Figure 3a,b). At 24 hpi, local NHP-treated leaves exhibited a significant increase in DAB staining intensity relative to uninfected baseline controls compared to mock- treated plants (∼25.5% vs. ∼15.7%, *p* < 0.001) (Figure 3c). This disparity expanded significantly by 48 hpi; while the ROS burst in mock-treated leaves diminished (∼13.7%), NHP-treated local leaves maintained a heightened oxidative state (∼34.3% intensity increase, *p* < 0.0001) (Figure 3c).

In distal leaves, the relative DAB staining intensity was initially comparable between NHP- and mock-treated plants at 24 hpi (∼15.6% vs. ∼12.9%) (Figure 3d). However, by 48 hpi, distal leaves of NHP-treated plants exhibited a significantly elevated and widespread H_2_O_2_ accumulation (∼18.7% increase) compared to waning levels in mock-treated distal leaves (∼5.8%; *p* < 0.001) (Figure 3d). These results demonstrate that localized NHP application primes a sustained local ROS burst and drives a delayed, systemic ROS wave in distal tissues during early *Pt* pathogenesis (Figure 3).

### Spatially separated NHP treatments differentiate early transcriptional reprogramming

To systematically dissect the spatial and temporal orchestration of the NHP-induced resistance (NIR) program, we conducted global transcriptomic profiling via RNA-sequencing (RNA-seq). Because SAR involves a highly coordinated signaling cascade that migrates from the site of initial perception to uninfected tissues, capture of this dynamic process requires examining both local (treated) and distal (untreated) leaves across critical earlier phases of fungal pathogenesis. We selected 24 and 48 hours post-infection (hpi) as our key time-points to catch the transition from early cellular defense priming to full systemic immune execution coinciding with systemic ROS signaling. By directly comparing uninfected controls (control), baseline compatible infections (mock+Pt), and NHP-treated and infected states (NHP+Pt), we aimed to isolate the precise transcriptional networks uniquely driven by NHP from those induced by normal pathogen challenge (Supplementary Figure 2).

Principal component analysis (PCA) of variance-stabilized RNA-seq counts revealed that tissue identity (local vs. distal) was the primary driver of transcriptional variation (PC1, 34%; Supplementary Figure 3a); notably, this substantial local vs. distal variance likely reflects a combination of inherent tissue-specific expression profiles and the localized physiological impact of syringe infiltration introducing mechanical stimuli to the treated leaf. While these localized treatment effects cannot be entirely decoupled from baseline tissue identity, the clear partition along the PC1 axis confirms that the localized physiological reaction remains structurally distinct from the systemic response of distal tissue. Concurrently, the treatment condition (control vs. mock+*Pt* vs. NHP+*Pt*) constituted the secondary driver of variation (PC2, 18%; Supplementary Figure 3b), indicating that NHP application exerts a profound regulatory influence that overrides standard host-pathogen interactions. This is further supported by an unsupervised bi-cluster heatmap, which demonstrated that NHP-treated samples formed a distinct transcriptional cluster entirely separate from normal *Pt* infection (mock+Pt) and uninfected controls (Supplementary Figure 3c).

Differential gene expression (DEG) analysis confirmed that local NHP treatment uniquely and systemically remodels the transcriptome. A direct comparison between the NHP+Pt and mock+Pt conditions, which was expected to reveal the specific gene networks uniquely engaged by NHP to actively restrict *Pt* colonization and override the host’s default susceptibility program, identified 2,274 DEGs. Nearly 80% (1,818) of these transcripts were up-regulated, indicating that NHP primarily promotes global gene expression induction rather than repression during early *Pt* pathogenesis (Supplementary Figure 4b). Of these, 616 genes (11.1%) were exclusively differentially expressed due to NHP addition during infection and were not significantly altered in baseline uninfected states (Supplementary Figure 4a). Hierarchical clustering of these 2,274 DEGs verified distinct, highly coordinated expression patterns across treatments and time points, establishing that NHP treatment leaves a massive systemic footprint that sets up an immune-ready state (Supplementary Figure 4c).

### NHP systemically induces genes involved in immune perception, redox regulation, and vesicle trafficking

NHP treatment promotes a coordinated immune-ready state rather than isolated defense outputs. Gene set enrichment analyses (GSEA) of the NHP-responsive transcriptome revealed a massive, systemic up-regulation of pathways governing pathogen perception and signal transduction. Transcripts encoding cell-surface receptor-like kinases (RLKs), notably cysteine- rich kinases (CRKs) and wall-associated kinases (WAKs) as well as intracellular immune receptors (NLRs), were heavily enriched (Figure 4a; Supplementary Figure 5).

**Figure 4.**
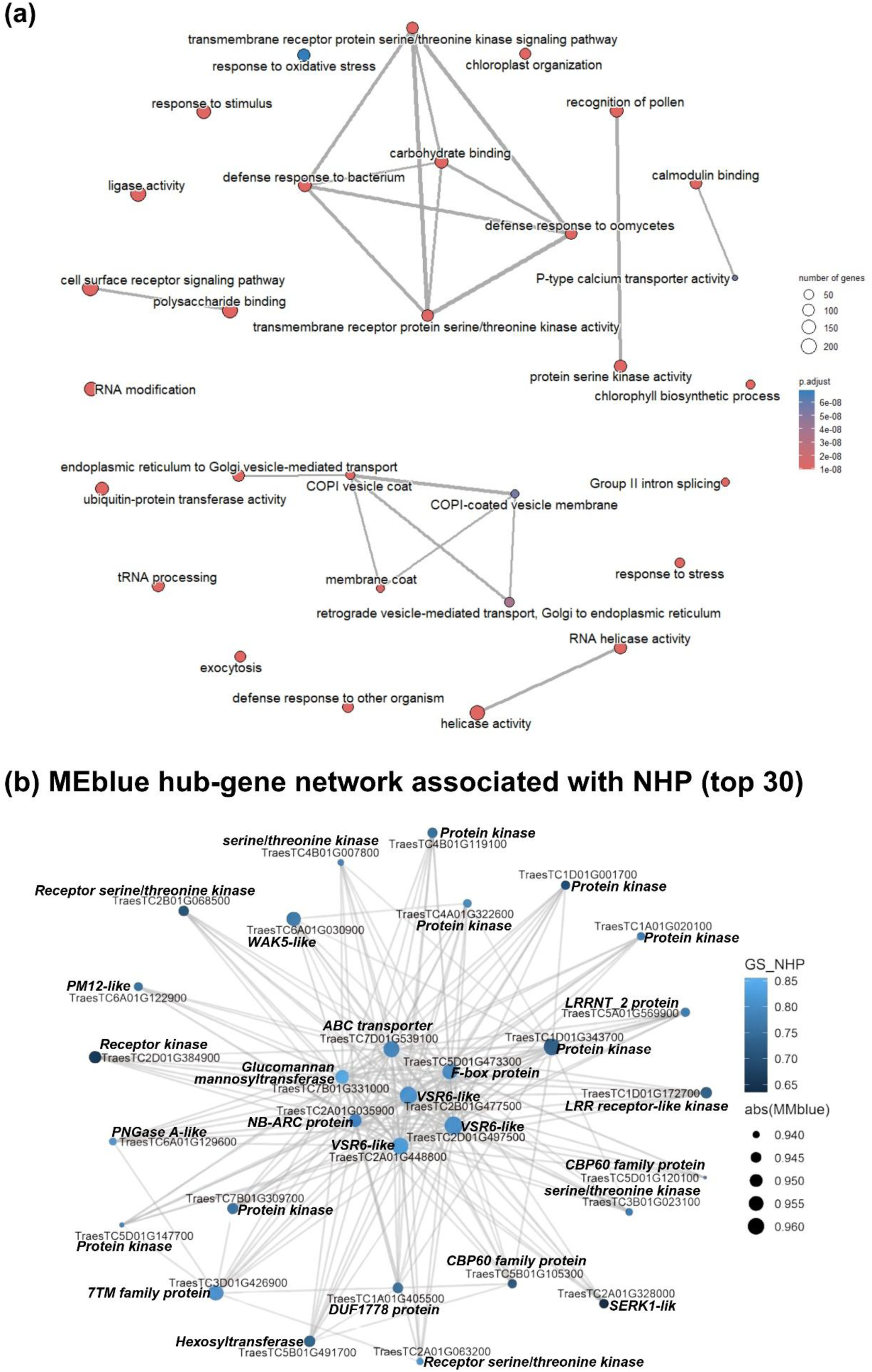
NHP treatment establishes a coordinated systemic immune-ready state and co- expression network. **(a)** Network plot of significantly enriched immune-related gene sets identified via Gene Set Enrichment Analysis (GSEA) comparing NHP+*Pt* vs mock+*Pt* conditions. Nodes (circles) represent distinct gene sets, with node size corresponding to the number of genes per set, and color indicating the adjusted *p*-value of enrichment significance. Lines connecting nodes represent significant functional interactions and shared gene membership. **(b)** Co-expression network of the top 30 hub genes within the highly NHP- responsive MEblue module identified via weighted gene co-expression network analysis (WGCNA). Node size reflects absolute module membership (|k_ME_|), and node color corresponds to gene significance (GS) for NHP treatment. Central hubs connecting the network include key PTI/ETI regulators (e.g., *WAK5*), vesicle trafficking components, and SAR-linked regulators. **Alt text:** Network diagrams highlighting enriched GSEA immune gene sets and top 30 hub genes in the NHP-responsive WGCNA MEblue co-expression module.

Downstream of pathogen perception, NHP treatment amplified signaling pathways essential for immune execution. Processes related to calcium influx, ER–Golgi vesicle trafficking (including SNARE-mediated endo- and exocytosis), and post-translational protein modifications (ubiquitination, phosphorylation) were significantly potentiated (Supplementary Figure 6). Weighted gene co-expression network analysis (WGCNA) further illuminated this coordinated transcriptional architecture, identifying a highly NHP-responsive co-expression module (MEblue; Supplementary Figures 7 and 8). Within this network, key PTI/ETI regulators (such as *WAK5*), vacuolar sorting receptors, and broad-spectrum resistance components act as highly connected hub genes orchestrating the NHP defense response (Figure 4b).

Corresponding to the whole-plant H_2_O_2_ burst observed phenotypically, NHP treatment robustly enriched pro-oxidant signaling pathways (including *SAG39* and cytochrome P450s) (Supplementary Figure 6a,b) while concurrently suppressing genes involved in bulk ROS- scavenging and antioxidation (Supplementary Figure 6c). NHP pre-treatment significantly down- regulated primary cellular metabolism, repressing transcripts linked to photosynthesis, sugar synthesis, lipid transport, and abiotic stress tolerance (Supplementary Figure 6d), which reflects energy trade-offs required to maintain this heightened immune activation. Thus, NHP treatment reallocates global plant resources toward defense and oxidative containment during early *Pt* pathogenesis.

### NHP treatment primes PTI, ETI, and broad-spectrum resistance gene networks

To identify gene candidates uniquely contributing to NHP-induced resistance, we filtered differentially expressed genes (DEGs) across temporal and spatial planes. NHP treatment exclusively up-regulated 1,679 genes during early *Pt* pathogenesis relative to both a baseline *Pt* infection and uninfected control states (Supplementary Figure 9a,b).

Cell-surface pathogen perception (PTI) was strongly enhanced by NHP, evidenced by the robust systemic induction of cysteine-rich receptor-like kinases (CRK10/25/40s) and wall- associated kinases (WAK1/2/3/5), alongside the induction of *Xa21*- and *FLS2*-like homologs (Figure 5; Table 1). Concurrently, intracellular effector-triggered immunity (ETI) was primed through the systemic induction of nucleotide-binding leucine-rich repeat receptors (NLRs), including *RPM1*-, *RGA3*-, *RPP13*-, and *Pik-1*-like genes (Figure 5; Table 1). NHP also potentiated the expression of durable, broad-spectrum resistance (*R*) genes; the functional allele of the ABC transporter *Lr34* on chromosome 7D (*Lr34res*) was systemically boosted at 24 hpi, alongside powdery mildew resistance orthologs *Pm12* and *Pm60* (Figure 5; Table 1).

**Figure 5.**
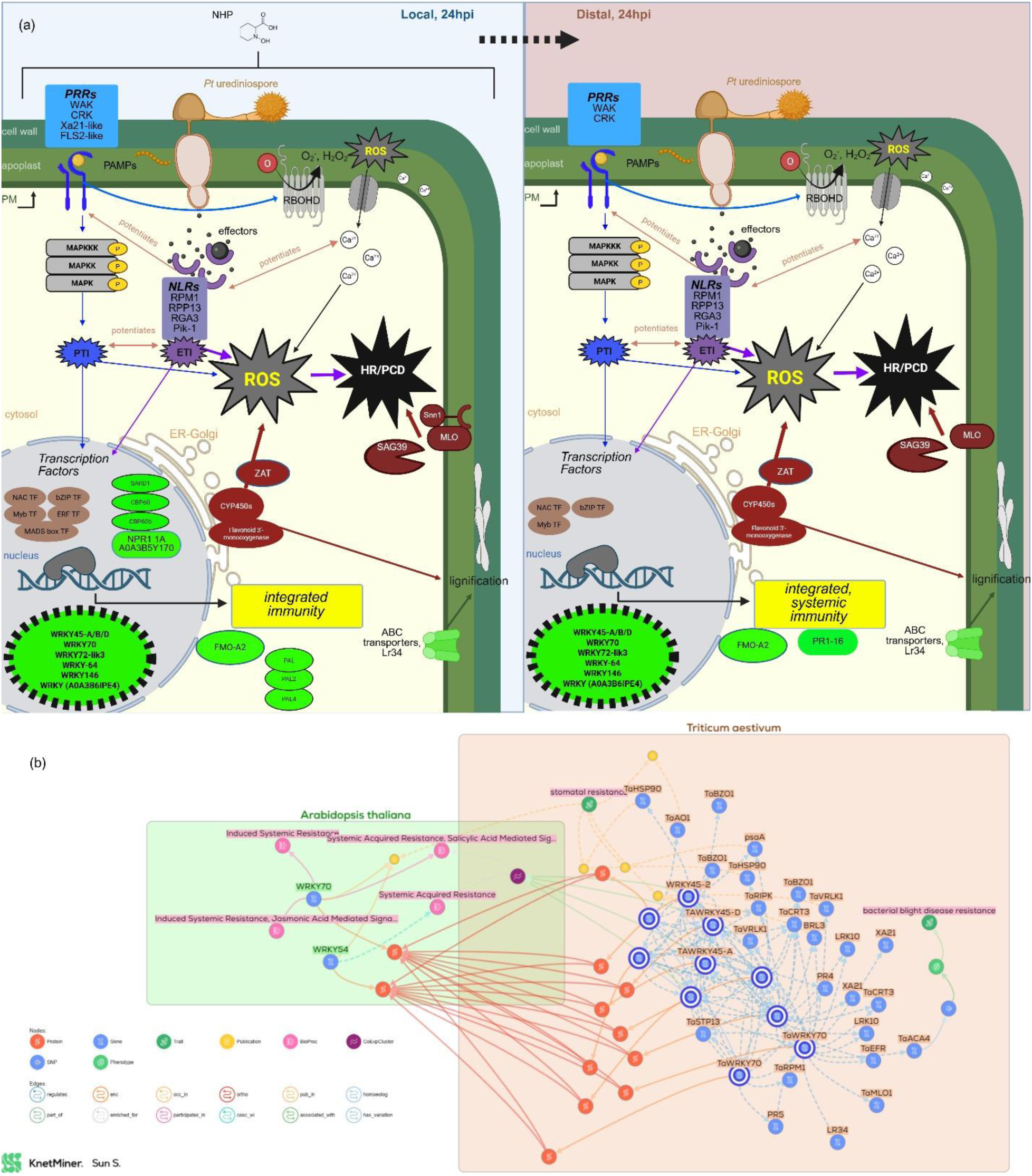
NHP primes local and systemic immune signaling networks associated with extensive WRKY induction. (a) Schematic model of enhanced pathogen perception and downstream immune signaling at 24 hours post-infection (hpi) in local leaves (left) and systemically primed distal leaves (right). NHP potentiates pattern-triggered immunity (PTI; blue) through plasma membrane-localized PRRs including WAKs, CRKs, *Xa21*-like, and *FLS2*-like receptors, alongside effector-triggered immunity (ETI; purple) via cytosolic NLRs such as *RPM1*, *RPP13*, *RGA3*, and *Pik-1*. These pathways converge on MAPK cascades, Ca²⁺ influx, and ROS production. Canonical SAR-associated regulators (green), including *NPR1*-like and *CBP60/SARD1* modules, act alongside broad-spectrum resistance transporters (e.g., *Lr34*) and pathogenesis-related genes (e.g., *PR1-16*) to establish integrated whole-plant immunity. **(b)** KnetMiner knowledge graph (Hassani-Pak et al., 2021) detailing the highly resolved functional interaction network of the top systemic WRKY transcription factors induced by NHP. The core subset of systemic WRKYs (including *TaWRKY45* and *TaWRKY70*) forms a highly connected network associated with multi-pathogen defense responses. The network graph can be viewed for more information: app.knetminer.com/network/7d5961f2-0329-47df-b477-1c57acd0e427. **Alt text:** Pathway schematic of local and systemic immune signaling convergence alongside a KnetMiner knowledge graph of systemic WRKY transcription factors.

**Table 1.** Representative NHP-responsive genes associated with PTI, ETI, and broad-spectrum resistance during early *Pt* infection.

| Immune layer | Gene / homolog | Thatcher gene ID | 24L | 24D | 48L | 48D | Functional / resistance evidence |
|---|---|---|---|---|---|---|---|
| **PTI** | *WAK5* | TraesTC6A01G030900 | 3.6 | 3.4 | — | — | NHP-associated MEblue top-30 hub |
|  |  | TraesTC3A01G555300 | 2.8 | 3.4 | — | — | — |
|  | *WAK2* | TraesTC6D01G036800 | 3.5 | — | — | — | Leaf-rust seedling and adult-plant resistance (Genievskaya et al., 2020; Mourad et al., 2024) |
|  | *CRK10* | TraesTC3D01G121300 | — | 9.7 | — | — | Annotated TaCRK70_3D (H. Liu et al., 2024) |
|  |  | TraesTC7A01G102700 | 2.7 | 2.9 | — | — | — |
|  | *CRK25* | TraesTC7A01G102800 | — | 2.4 | — | — | TaCRK3; sharp eyespot/FCR resistance-associated (Guo et al., 2021; Q. Li et al., 2024; H. Liu et al., 2024) |
|  |  | TraesTC7B01G077900 | — | 7.7 | — | — | — |
|  |  | TraesTC5D01G622200 | 4.5 | 3.9 | — | — | — |
|  | *CRK40* | TraesTC4D01G101700 | 6 | 4.5 | — | — | — |
|  | *XA21-like* | TraesTC4D01G409400 | 5.9 | — | — | — | Homolog of rice XA21; LRR-RLK XII-a<br>(Cantu et al., 2013; S. Liu et al., 2023) |
|  | *FLS2-like* | TraesTC6D01G409200 | 6.8 | — | — | — | Flagellin receptor |
| **ETI** | *RPP13-like* | TraesTC2D01G005900 | 2.8 | — | — | — | Candidate in a high-confidence meta-QTL associated with resistance to leaf rust, stripe rust, powdery mildew, and Septoria tritici blotch (Amo & Soriano, 2022) |
|  | *RGA3-like* | TraesTC2D01G673200 | — | 3.5 | — | — | *P. tritici-repentis* defense/cell-death associated (Moolhuijzen et al., 2023) |
|  | *Pik-1-like* | TraesTC2B01G374400 | 7 | — | — | — | — |
|  | *RPM1-like* | TraesTC7D01G520900 | 4.2 | 3 | — | — | — |
|  |  | TraesTC7B01G403100 | 4.3 | 3.3 | — | — | — |
|  |  | TraesTC2B01G664000 | 5.2 | 5.1 | — | — | — |
|  |  | TraesTC1D01G025600 | 2.8 | — | 2.7 | — | Putative NBS-LRR disease-resistance gene within the *Lr42* leaf-rust resistance interval (Y. Liu et al., 2021) |
|  | *Lr34-7D* | TraesTC7D01G097700 | 3.1 | 2.5 | — | — | Durable broad-spectrum resistance ABC transporter |
| **Broad-spectrum** | *Pm12-like†* | TraesTC6A01G122900 | — | — | — | — | Powdery mildew resistance NLR; HR-associated resistance; NHP-associated MEblue top-30 hub |
|  | *Pm60†* | TraesTC7B01G522900 | — | — | — | — | Powdery mildew resistance NLR; HR-associated resistance |

L2FC, log2 fold change; 24L and 48L, local NHP-treated leaf tissue collected at 24 and 48 hours post-infection (hpi), respectively; 24D and 48D, untreated distal leaf tissue collected at the corresponding time points. Values represent L2FC in NHP-treated, *P. triticina*-infected plants (NHP+Pt) relative to mock-treated, *P. triticina*-infected plants (mock+Pt) for the indicated tissue and time point. Positive values indicate higher transcript abundance following NHP treatment. A dash (–) indicates that the gene did not meet the stringent differential-expression threshold used for the tissue- and time-specific candidate analysis.†*Pm12*-like and *Pm60* were significantly upregulated in the overall NHP+Pt versus mock+Pt comparison (L2FC = 1.53 and 1.74; adjusted *P* = 5.75 × 10<sup>−8</sup> and 7.49 × 10<sup>−7</sup>, respectively), but did not meet the more stringent L2FC > 2 threshold applied to the individual tissue- and time-specific candidate analysis.

These widespread defense responses were accompanied by the NHP-specific induction of over 50 WRKY transcription factors. A core systemic subset – including all three *TaWRKY45* homeologs, *TaWRKY70*, and *TaWRKY64* – was strongly induced locally at 24 hpi and remained highly expressed in distal leaves through 48 hpi (Figure 5; Table 2). Furthermore, NHP treatment activated canonical SAR-associated regulatory nodes, including a previously uncharacterized *NPR1*-like triad (chromosomes 1A/B/D), *TaSARD1* and *CBP60* family members, multiple *PAL* genes (*PAL2*, *PAL4*), and downstream pathogenesis-related transcripts such as *PR1-16* (Figure 5; Table 2). Together, these data show that NHP treatment induces a multi-layered systemic immune state encompassing receptor, transcriptional, and broad-spectrum resistance gene networks in hexaploid wheat.

**Table 2.** Representative NHP-responsive transcriptional regulators and systemic acquired resistance-associated genes during early *Puccinia triticina* infection.

| Regulatory layer | Gene / homolog | Thatcher gene ID | NHP-responsive expression | Functional annotation / prior resistance evidence |
|---|---|---|---|---|
| **Systemic WRKY regulation** | *TaWRKY45-A* | TraesTC2A01G512800 | 24L (2.8) / 24D (3.0) / 48D (3.7) | SA-inducible; overexpression enhances powdery mildew and leaf-rust resistance independently of *Pm3* and *Lr34* (Bahrini, Ogawa, et al., 2011b); implicated in stripe-rust interactions (Yao et al., 2025) |
|  | *TaWRKY45-B* | TraesTC2B01G552200 | 24L (2.6) / 24D (2.9) / 48D (2.9) | SA-responsive; associated with *Fusarium graminearum* and *Xanthomonas translucens* responses (Heckmann et al., 2024b) |
|  | *TaWRKY45-D* | TraesTC2D01G570500 | 24L (3.5) / 24D (4.1) / 48D (2.7) | BTH-responsive and associated with FHB resistance bahrini 2011, hu 2019 (Bahrini, Ogawa, et al., 2011b; Hu et al., 2019b) |
|  | *WRKY70* | TraesTC2B01G130600 | 24L (2.4) / 24D (3.1) / 48D (4.1) | Associated with induced systemic resistance and *Pyrenophora tritici-repentis* responses (Andersen et al., 2021; L. Li et al., 2022b) |
|  | *WRKY64* | TraesTC4D01G207900 | 24L (5.7) / 24D (5.6) / 48D (2.4) | Associated with *X. translucens* responses (Shah et al., 2023b), FHB resistance (Walker et al., 2024b), and all-stage stripe-rust resistance (Yang et al., 2023b) |
| **SAR regulation** | *NPR1/NIM1-like* | TraesTC1A01G247600 | 24L (5.8) | Previously uncharacterized NPR1-like wheat candidate |
|  |  | TraesTC1B01G273400 | – | Previously uncharacterized NPR1-like wheat candidate |
|  |  | TraesTC1D01G268900 | – | Previously uncharacterized NPR1-like wheat candidate |
|  | *SARD1-like* | TraesTC3A01G036000 | 24L (3.4) | Pathogen-responsive candidate |
|  |  | TraesTC3B01G057000 | 24L (3.7) | Associated with multiple pathogen-resistance QTL (Saini et al., 2022) |
|  |  | TraesTC3D01G048800 | 24L (4.3) | Associated with stripe-rust/FHB resistance in an alien chromosome-substitution line (Du et al., 2022) |
|  | *CBP60-like* | TraesTC5B01G105300 | 24L (2.8) | NHP-associated MEblue top-30 hub gene |
| **SA-associated metabolism** | *PAL4* | TraesTC2B01G425900 | 24L (3.7) | NHP-promoted PAL; previously associated with stress/pathogen responses (Jasrotia et al., 2022; Mathew et al., 2019; Rasool et al., 2021; Su et al., 2023; Yue et al., 2021; Y. Zhang et al., 2019a) and *Lr34*-associated defense (Y. Zhang et al., 2024b) |
|  | *PAL2* | TraesTC2D01G447600 | 24L (3.3) | NHP-induced PAL associated with wheat stress/pathogen responses (Davoudnia et al., 2024; Kroll et al., 2025; L. Li et al., 2023; Vaughan et al., 2022; M. Zhang et al., 2021; Y. Zhang et al., 2019b) |
| **Defense output** | *PR1-16* | TraesTC5D01G502100 | 24D (5.6) | Pathogen-responsive PR1 associated with *P. tritici-repentis*, FHB, beneficial microbial inoculation, and *X. translucens* responses |

L2FC, log2 fold change; 24L and 48L, local NHP-treated leaf tissue collected at 24 and 48 hours post-infection (hpi), respectively; 24D and 48D, untreated distal leaf tissue collected at the corresponding time points. Values represent L2FC in NHP-treated, *P. triticina*-infected plants (NHP+Pt) relative to mock-treated, *Pt*-infected plants (mock+Pt) for the indicated tissue and time point. Positive values indicate higher transcript abundance following NHP treatment. A dash (–) indicates that the gene did not meet the stringent differential-expression threshold used for the tissue- and time-specific candidate analysis.

### NHP treatment induces systemic accumulation of SA, SAG, Pip, and NHP without suppressing JA

To anchor our transcriptional observations in corresponding chemical changes, we utilized targeted LC-MS to quantify relative levels of the important SAR-signaling molecules and pathogen defense-associated phytohormones like Pip, NHP, SA, SA glucoside (SAG), and jasmonic acid (JA) during early *Pt* pathogenesis. Metabolites were quantified using authentic external standards and normalized to sample fresh weight; therefore, interpretation focused primarily on treatment-dependent differences within the experiment. Metabolite levels were measured in local treated leaves and distal untreated leaves at 24 and 48 hpi across uninfected controls (Figure 6a), mock water-treated plants or inoculated with *Pt* (Figure 6b), and NHP- treated plants inoculated with *Pt* (Figure 6c).

**Figure 6.**
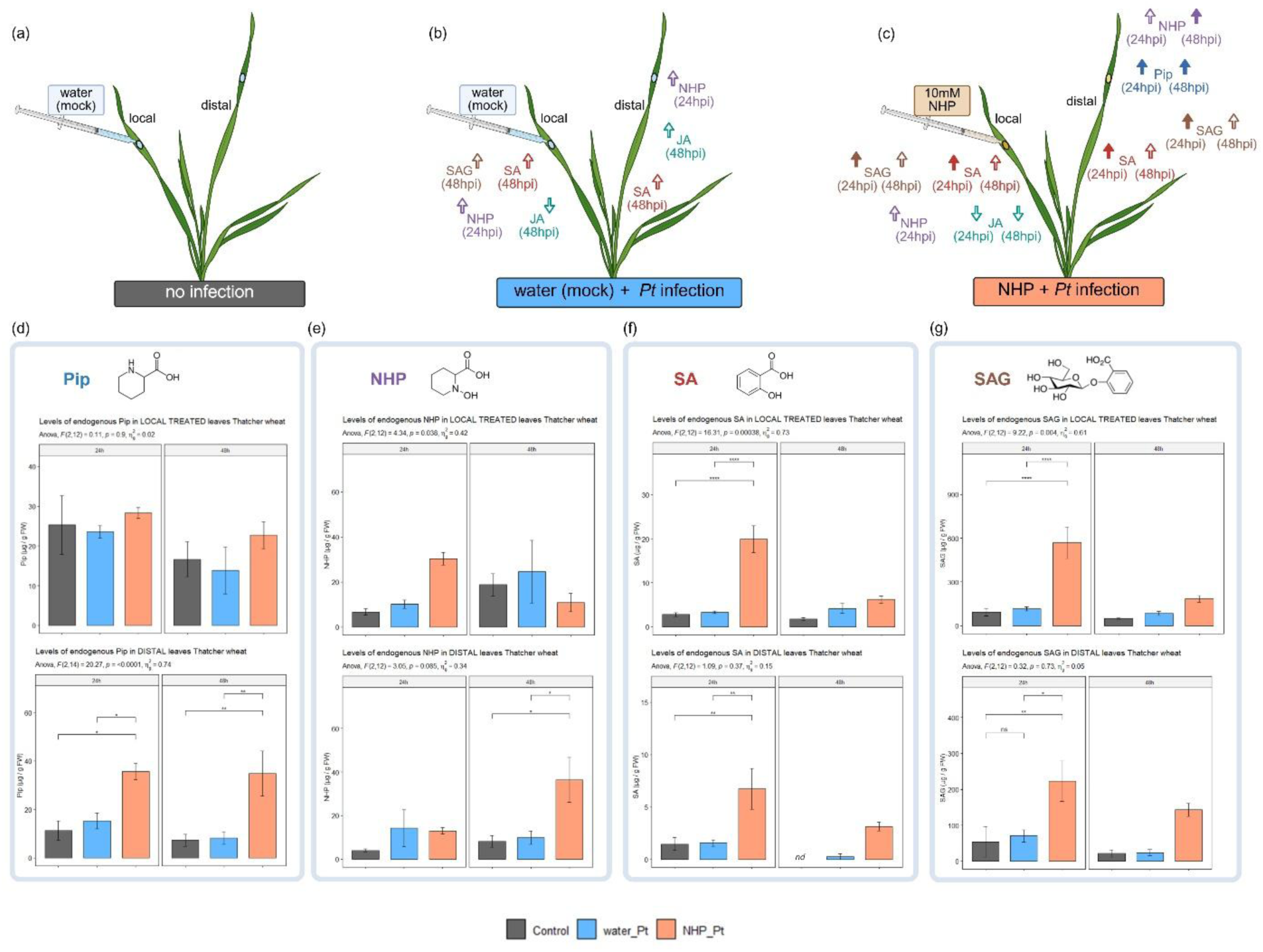
Local NHP supplementation leads to significantly different accumulation of Pip, NHP, SA, and SAG (but not JA) across treatments and time points. Four-leaf stage Thatcher wheat plants were infiltrated in the second true leaf with either a mock treatment (water) or 10 mM NHP. At 24 hours post-treatment, plants were either systemically inoculated with *Pt* race 1 BBBD urediniospores or left uninfected (control). Local treated leaves and distal untreated leaves were harvested at 24 and 48 hours post-infection (hpi) for targeted LC-MS analysis. (a–c) Schematic summary of overall metabolite accumulation patterns for uninfected controls (a), mock-treated plants with Pt infection (b), and NHP-treated plants with Pt infection (c). Filled arrows in (b,c) summarize statistically significant changes in metabolites relative to uninfected controls; unfilled arrows denote visible trends where standard errors do not overlap but differences lack statistical significance via ANOVA. (d–g) Quantitative LC-MS measurements of endogenous Pip (d), NHP (e), SA (f), and SAG (g). Bar plots display concentrations (μg/g fresh weight) for local treated leaves (top row) and distal untreated leaves (bottom row) at 24 and 48 hpi. Grey bars represent uninfected controls, blue bars represent mock-treated infected plants, and orange bars represent NHP-treated infected plants. Graph data are presented as mean values ± SEM from three independent experiments (*n* = 3 biological replicates), with each replicate pooling tissue from five individual plants. A two-way ANOVA (Type II test) with Bonferroni post-hoc pairwise comparisons were conducted (* *p* < 0.05, ** *p* < 0.01, *** *p* < 0.001, **** *p* < 0.0001; *nd* = not detected; blank = not significant. **Alt text:** LC-MS bar plots showing local and systemic accumulation of Pip, NHP, SA, and SAG at 24 and 48 hours post-*Pt* infection following local NHP treatment.

NHP treatment did not significantly alter Pip accumulation in local leaves during *Pt* infection; however, it significantly increased Pip levels in distal, untreated leaves (Figure 6d). In NHP-treated plants, distal Pip reached ∼35.7 μg/g FW at 24 hpi and ∼34.8 μg/g FW at 48 hpi, representing significant ∼2-fold and ∼4-fold increases relative to a baseline *Pt* infection, respectively (Figure 6d). Local NHP treatment also altered systemic NHP distribution patterns; while NHP levels in distal leaves remained similar across all treatments at 24 hpi, they rose significantly in NHP-treated plants by 48 hpi (∼36.3 μg/g FW vs ∼8–10 μg/g FW in control and mock-treated plants) (Figure 6e). This ∼4-fold systemic NHP increase coincides with the 48 hpi surge of distal H_2_O_2_ and the consequent global transcriptional activation of ROS-regulating defense genes.

Both SA and its storage form, SAG, were strongly elevated by NHP at 24 hpi in both local and distal leaves (Figure 6f, g). *Pt* infection alone was not sufficient to significantly alter SA or SAG throughout systemic foliar tissue at these early time-points (Figure 6f, g). However, in local leaves at 24 hpi, NHP-treated plants showed a ∼6-fold higher accumulation of SA (∼19.9 μg/g FW) (Figure 6f) and a ∼5-fold higher accumulation of SAG (∼568.8 μg/g FW) (Figure 6g) compared to mock controls. Distal leaves displayed a similar pattern at 24 hpi, with ∼4-fold SA (Figure 6f) and ∼3-fold SAG (Figure 6g) increases. At 48 hpi, SA and SAG levels remained elevated systemically in NHP-treated plants relative to controls, though differences were no longer statistically significant due to measurement variability (Figure 6f, g). Notably, SAG levels were consistently observed at approximately 30-fold higher concentrations relative to free SA (Figure 6f, g).

In contrast to the SA and NHP pathways, JA accumulation was not significantly affected by NHP treatment across any tissue or time point (Figure 6a–c; Supplementary Figure 10). While baseline *Pt* infection consistently showed slightly higher distal JA relative to both uninfected controls and NHP-treated plants, these differences did not reach statistical significance. Thus, NHP treatment does not appear to significantly antagonize JA production during early *Pt* infection, highlighting a crucial divergence from the canonical hormone crosstalk observed in eudicot SAR models.

### Wheat *FMO* candidates respond to NHP treatment in a clade-specific manner

Given the systemic NHP accumulation observed following NHP treatment and *Pt* infection, we next examined the wheat genes associated with NHP biosynthesis and their transcriptional responses under these conditions. To identify wheat orthologs of the Arabidopsis NHP-biosynthetic enzymes, we mapped the previously identified 170 *TaFMO* genes from the wheat cv. Chinese Spring reference genome (Sun and Bakkeren, 2024) onto the cv. Thatcher genome, identifying 163 *TaTC-FMO* candidates. Phylogenetic clustering partitioned these candidates into three major clades A, B, and C (Supplementary Figure 11). In contrast to Arabidopsis where *AtFMO1* is a singular indispensable gene for NHP synthesis, only one *TaTC- FMO* (*TraesTC4B01G422800*) from sub-clade A2 was significantly up-regulated systemically by NHP treatment during *Pt* infection. Two additional ‘non-canonical’ *TaTC-FMOs* from sub- clade B2 were differentially expressed but responded primarily to a baseline *Pt* infection rather than NHP specifically (Supplementary Figure 11). This suggests an extensive FMO redundancy and divergent pathogen-responsive specialization in hexaploid wheat.

### Partial silencing of *TaSARD4* does not produce a detectable change in *Pt* resistance

In contrast to the highly expanded *FMO* family, *SARD4* orthologs were highly conserved as single-copy genes across diploid angiosperms, with three homeologs (A, B, and D) identified in wheat grouping tightly with other monocot species (Figure 7a). All three *TaSARD4* homeologs were constitutively expressed in local and distal tissues, but were not significantly induced by *Pt* infection or NHP treatment. To investigate the functional relevance of *TaSARD4 in planta*, we simultaneously silenced all three homeologs using a barley stripe mosaic virus (BSMV)-induced gene silencing (VIGS) construct targeting a conserved 401-bp fragment (Supplementary Figure 12; Supplemental Methods S3). While the BSMV-VIGS treatment successfully achieved a ∼5.7- fold reduction in *TaSARD4* transcripts at 24 hpi relative to the empty vector control (Figure 7b,c), this knockdown did not significantly alter *Pt* infection frequency on systemic leaves of the VIGS-treated plant (Figure 7d). However, because this VIGS approach resulted in a partial transient knockdown rather than a complete knockout, it is possible that residual *TaSARD4* activity, or alternative redundant biosynthetic pathways, are sufficient to maintain baseline resistance during early *Pt* pathogenesis.

**Figure 7.**
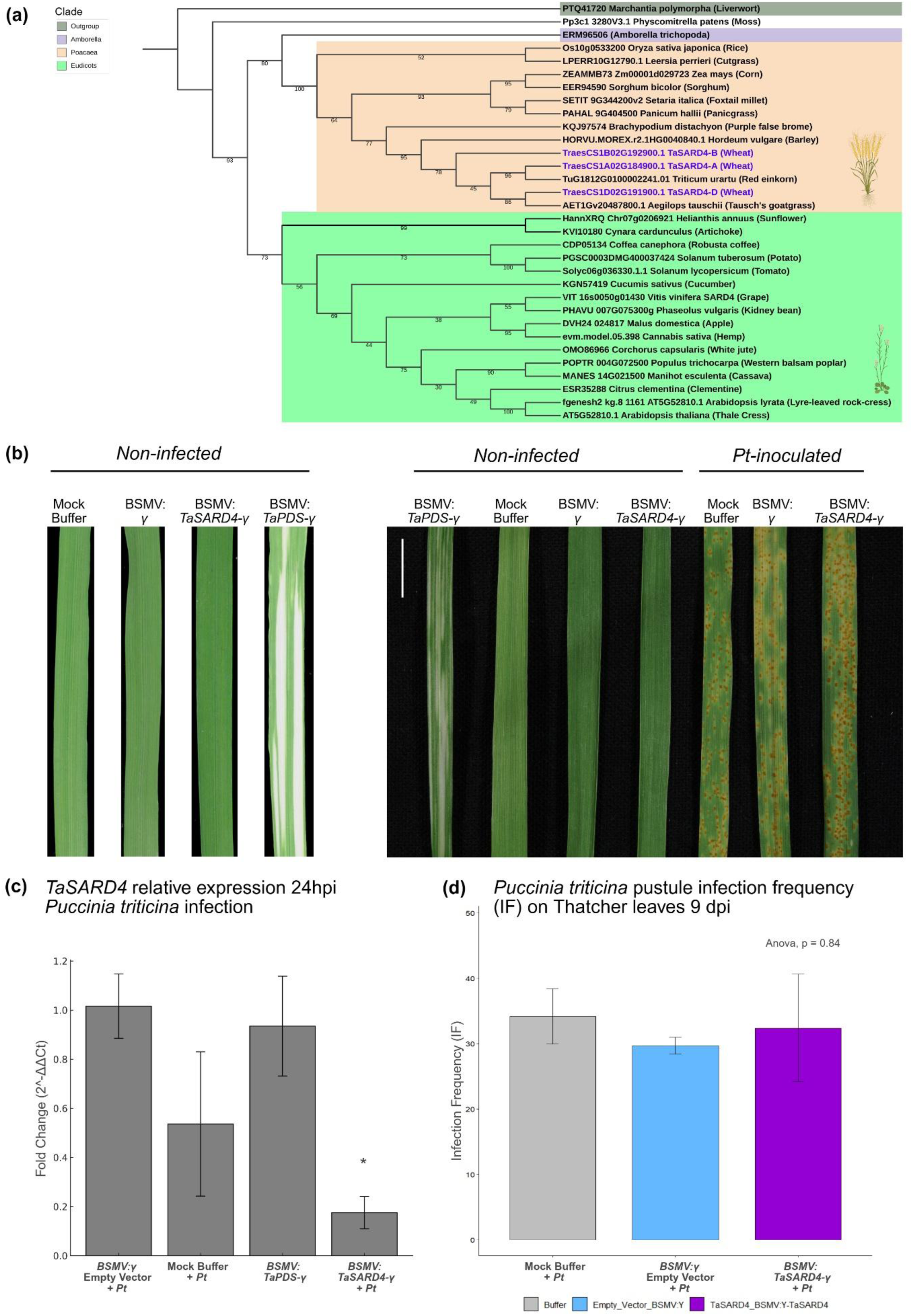
Evolutionary conservation and transient silencing of *TaSARD4* homeologs during *Pt* infection. **(a)** Maximum-likelihood phylogenetic tree of SARD4 protein sequences across 30 diverse plant species (IQ-TREE, 1,000 bootstrap replicates, JTT+G4 model). The wheat (*Triticum aestivum* cv. Thatcher) *TaSARD4-A*, *-B*, and *-D* homeologs (purple text) group distinctly within the monocot clade (orange), separated from eudicots including *Arabidopsis thaliana* (green). **(b)** Representative line-up of the systemic fourth leaf from uninfected and *Pt*- inoculated plants treated with BSMV-VIGS constructs or a mock buffer control at 9 days post- inoculation (dpi). Scale bar = 1 cm. **(c)** Relative *TaSARD4* mRNA expression in the systemic fourth leaf at 24 hours post-*Pt* infection. Bars represent the fold-change (2^-ΔΔCt) of *TaSARD4* transcripts in plants inoculated with empty vector (BSMV:γ), mock buffer, PDS silencing vector (BSMV:TaPDS-γ), and the *TaSARD4* silencing vector (BSMV:TaSARD4-γ). Asterisk indicates a significant ∼5.7-fold reduction compared to empty vector (Welch’s t-test, *p* = 0.011). **(d)** Average *Pt* infection frequency (IF; pustules per cm²) on the systemic fourth leaf at 9 dpi for buffer-treated (grey), empty vector (blue), and BSMV-γ-TaSARD4 silenced plants (purple). Bars represent means ± SEM of three independent biological replicates (n=3; each comprising six individual plants). Differences were not statistically significant (ANOVA, *p* = 0.84). **Alt text:** Phylogenetic tree, leaf photos, and bar plots showing monocot *TaSARD4* homeolog conservation, VIGS transcript silencing (∼5.7-fold), and *Pt* infection frequency.

To test whether the *TaSARD4* enzymes possess conserved biochemical functionality, we constitutively expressed the Thatcher *TaSARD4-A*, *-B*, and *-D* homeologs in an Arabidopsis *Atsard4-KO* deletion line deficient in NHP biosynthesis (Figure 8a, b). To synchronize immune induction, transgenic seedlings were elicited with the pattern-triggered immunity (PTI) elicitor NLP20 (Böhm et al., 2014), and the stable storage sink of NHP, glycosylated NHP (NHP-*O*Glc), was measured. NLP20 elicitation successfully induced NHP-*O*Glc accumulation in wild-type Columbia-0 plants, which was abolished in the *Atsard4-KO* line. Notably, transgenic lines overexpressing *TaSARD4-B* (lines B1, B2) successfully restored NLP20-triggered NHP-*O*Glc accumulation to near wild-type levels (Figure 8c). While *TaSARD4-A* and *-D* lines did not show significant restoration in this assay, the successful complementation by *TaSARD4-B* confirms that wheat retains a functional SARD4 enzyme capable of facilitating NHP biosynthesis, even if its *in planta* regulation diverges from the canonical Arabidopsis model.

**Figure 8.**
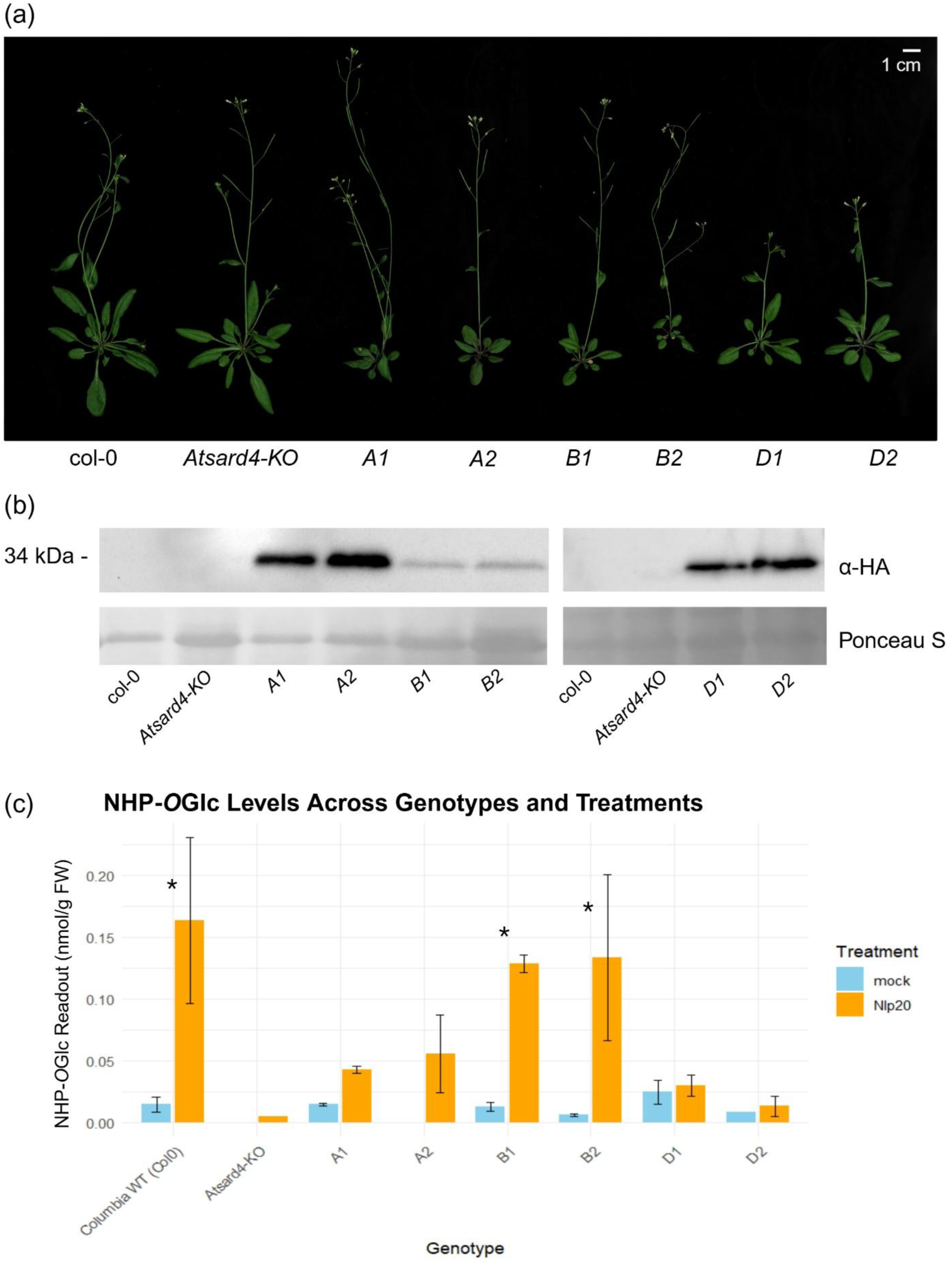
Heterologous expression of wheat *TaSARD4-B* restores NHP-*O*Glc accumulation in an Arabidopsis *sard4* knockout mutant. **(a)** Representative line-up of 5-week-old *A. thaliana* lines. Genotypes include Col-0 (wild-type), *Atsard4-KO*, and independent stable transgenic lines overexpressing Thatcher wheat homeologs: *TaSARD4-A* (A1, A2), *TaSARD4-B* (B1, B2), and *TaSARD4-D* (D1, D2). Scale bar = 1 cm. **(b)** Immunoblot analysis confirming *TaSARD4* protein expression in 10-day-old transgenic seedlings utilizing an anti-hemagglutinin (α-HA) antibody. Ponceau S staining served as a loading control. **(c)** Quantification of glycosylated NHP (NHP-*O*Glc) levels (nmol g⁻¹ fresh weight) in local leaves 24 hours after infiltration with mock buffer (blue) or the PTI-elicitor NLP20 (yellow). Bars represent means ± SEM of three independent biological replicates (*n* = 3, pooled from six plants each). Asterisks indicate significant induction between mock and NLP20 treatments within genotypes (ANOVA with Tukey HSD post-hoc test, *p* < 0.05). Overexpression of *TaSARD4-B* (lines B1, B2) successfully restored NLP20-induced NHP-*O*Glc accumulation to wild-type Col-0 levels. **Alt text:** Plant photographs, Western blot, and bar plot showing TaSARD4-B overexpression restores NLP20-induced NHP-*O*Glc production in Arabidopsis sard4 mutants.

## Discussion

In this study, we demonstrate that local application of NHP induces robust systemic immunity in hexaploid wheat against the leaf rust fungus *Puccinia triticina*. Exogenous NHP acts as a master immune amplifier, significantly restricting fungal colonization and triggering a coordinated systemic reactive oxygen species (ROS) burst. Integrating histological, transcriptomic, and metabolomic analyses, we reveal that this NHP-induced resistance (NIR) diverges from canonical eudicot models that operates via a synergistic Pip–NHP–SA metabolic axis lacking jasmonic acid antagonism, utilizes cereal-specific FMO regulatory architecture, and globally primes broad-spectrum resistance networks alongside extensive WRKY transcriptional reprogramming.

### NHP treatment elicits a coordinated, ROS-driven systemic immune state in wheat

ROS bursts are dominated by O_2_^-^ and H_2_O_2_ and form part of the early hypersensitive response (HR) that limits pathogen spread (Grant & Loake, 2000; Jan et al., 2025; Mittler, 2017). A biphasic ROS pattern characterizes plant–pathogen interactions, such that a transient early burst common to both compatible and incompatible interactions occurs at the first pathogen perception, followed by a stronger, sustained burst only in resistant (incompatible) interactions that promotes programmed cell death (PCD) (Grant & Loake, 2000; Hao et al., 2022). Localized NHP pre-treatment is sufficient to trigger a robust, SAR-like systemic immunity in hexaploid wheat against the obligate biotrophic fungal pathogen *Pt*. In typical resistant (incompatible) wheat interactions with rust fungi, an early and intense H_2_O_2_ accumulation at 12–24 hpi strongly correlates with resistance by restricting pathogen spread, whereas susceptible cultivars fail to establish this early ROS surge (Gou et al., 2015; Wang et al., 2007). Our findings demonstrate that exogenous NHP transforms a normally compatible Thatcher–*Pt* interaction into one more closely resembling an incompatible response, producing a pronounced H_2_O_2_ accumulation at 24– 48 hpi in local leaves that subsequently spreads to distal leaves by 48 hpi (Wang et al., 2007). Histological DAB visualization revealed that NHP treatment triggers ROS not only at stomatal penetration sites but also broadly across adjacent mesophyll cells, which is consistent with ROS propagation during early haustorial establishment (Wang et al., 2007). This amplified ROS environment is likely prohibitive toward early *Pt* colonization, explaining the observed systemic reduction in *Pt* fungal biomass and overall disease severity.

The biphasic accumulation of ROS requires precise coordination, and our transcriptomic data indicates that NHP fundamentally reshapes redox homeostasis to favor sustained oxidative signaling over bulk antioxidant scavenging (Mittler, 2017; Mittler et al., 2004). NHP treatment exclusively induced the transcription of a respiratory burst oxidase homolog-encoding gene (*RBOH*, *TraesTC1D01G323900*) during *Pt* infection, potentially driving the elevated local and systemic H_2_O_2_ levels, mirroring NADPH oxidases up-regulated during early pathogen-induced ROS amplification (Wu et al., 2023). Furthermore, NHP treatment strongly influenced transcriptional regulators modulating ROS dynamics, such as *WRKY40* and *WRKY19-a*. *WRKY40* expression was more strongly promoted by NHP than *WRKY19-a*, which likely biases the immune system toward sustained positive ROS production, consistent with the competitive dual regulatory module these two TFs exert on ROS production (He et al., 2025).

NHP treatment also significantly up-regulated the transcripts of several genes with established roles in hypersensitive response (HR) and programmed cell death (PCD) execution, including *SAG39* cysteine proteases (Liu et al., 2010; Wang et al., 2022), *ZAT11* zinc finger transcription factors (Qureshi et al., 2013), and prohibitin-domain proteins (PHBs) (Chen et al., 2005; Nadimpalli et al., 2000; Wen et al., 2014). Interestingly, NHP pre-treatment also induced susceptibility-linked cell-death regulators, such as an *MLO*-like gene (Bhat et al., 2005; Büschges et al., 1997; Jørgensen, 1992; Kusch et al., 2016) and the *Snn1* sensitivity gene (Kariyawasam et al., 2023; Liu et al., 2012; Shi et al., 2016). This suggests that NHP may co-opt and reshape existing susceptibility-linked PCD pathways to accelerate cell-death regulation during early infection.

Importantly, while NHP treatment promoted pro-oxidant and signaling-associated ROS components, it did not concurrently elevate bulk ROS-scavenging capacity. Genes encoding key enzymatic antioxidants, such as catalases (CATs), superoxide dismutase (SODs), and most peroxidases (PODs), which can sometimes increase susceptibility by prematurely reducing H₂O₂ (Prasad et al., 2019; Savadi et al., 2018), were not significantly induced by NHP treatment. Furthermore, pathways regulating non-enzymatic antioxidants, such as carotenoids (Jan et al., 2025), ascorbic acid (Viviani et al., 2021), and quercetin (Arikan et al., 2023), were enriched among the gene sets negatively affected by NHP. By suppressing these antioxidant scavenging pathways, NHP prolongs ROS retention, driving a sustained, heightened oxidative state that effectively contains the biotrophic rust fungus early in its infection cycle, consistent with previous findings that lowered antioxidants promote stronger immune outputs (Pavet et al., 2005).

### NHP treatment primes broad-spectrum resistance through integrated PTI, ETI, and WRKY networks

Rather than activating a single, isolated defense pathway, NHP functions as a global immune amplifier by coordinately priming both pattern-triggered immunity (PTI) and effector- triggered immunity (ETI) networks during early *Pt* pathogenesis. Cell-surface receptor-like kinases (RLKs), which perceive pathogen and damage-associated molecular patterns (MAMPs/DAMPs), are heavily modulated during wheat-rust interactions and contribute to long- distance immune signaling (Couto & Zipfel, 2016; Osakabe et al., 2013; Sharma et al., 2013; Shiu & Bleecker, 2001; Singh et al., 2022; Wan et al., 2019; Wang et al., 2019). We observed that NHP treatment systemically increased transcription of multiple RLK subclasses, notably cysteine-rich kinases (CRKs) and wall-associated kinases (WAKs), which govern ROS production, callose deposition, and downstream signal transduction (Anderson et al., 2001; K. Chen et al., 2004; He et al., 2025; Mapuranga et al., 2022; Saintenac et al., 2021; Verica & He, 2002; J. Wang et al., 2021; Q. Wu et al., 2024; Y. Zhang et al., 2023). Furthermore, intracellular nucleotide-binding leucine-rich repeat receptors (NLRs), which form the core sensors of ETI (Andersen et al., 2020; Duxbury et al., 2021; El Kasmi, 2021; Jones et al., 2016), were broadly potentiated, with RPM1, RPP13, RGA3, and Pik-1 orthologs systemically induced at 24 hpi. RPM1 orthologs trigger Ca^2+^ influx and HR (Bi et al., 2021; M. Grant et al., 2000; Xu et al., 2022), and a wheat TaRPM1 confers stripe rust resistance via SA-associated pathways (Wang et al., 2020). The early, systemic induction of these receptors establishes NHP as an inducer of broad-spectrum resistance acting at both the cell-surface and cytosolic interface.

The primed state established by NHP treatment also promotes the expression of durable, broad-spectrum resistance (*R*) genes extending beyond rust-specific effectors. Transcripts encoding the functional allele of the ABC transporter *Lr34* on chromosome 7D (*Lr34res*) were systemically boosted 24 hours after NHP treatment. *Lr34res* confers non-race-specific adult plant resistance against leaf rust, stripe rust, and powdery mildew by reinforcing cell walls via increased lignin precursors, modulating SA/JA accumulation, and altering phospholipid signaling linked to ROS production (Chauhan et al., 2015; Deppe et al., 2018; Krattinger et al., 2009, 2011, 2013, 2019; Lillemo et al., 2013; McCallum & Hiebert, 2022; Nakano et al., 2013; Risk et al., 2013; Silva et al., 2015; Zhang et al., 2024; Zhao, 2015; Zhao & Wang, 2013). Additionally, NHP treatment systemically induced powdery mildew resistance orthologs like *Pm12* and *Pm60*, which encode NLRs conferring HR-associated resistance to *Blumeria graminis f. sp. tritici* (An et al., 2019; M. Li et al., 2024; Niewoehner & Leath, 1998; Z. Wu et al., 2023a; D. Zhang et al., 2019; S. Zhu et al., 2023; Zou et al., 2018). This demonstrates that NHP can act upstream to amplify diverse, multigenic resistance hubs, further positioning it as a master amplifier of durable crop resistance.

To coordinate these widespread PTI, ETI, and broad-spectrum defense outputs, NHP relies heavily on the induction of WRKY transcription factor networks. Our transcriptome analysis revealed that NHP treatment exclusively induced over 50 WRKY TFs, with a core subset remaining highly induced systemically through 48 hpi. Notably, all three homeologs of *TaWRKY45* and *TaWRKY70* emerged as focal systemic integrators. In cereals, *WRKY45* orthologs are master regulators of benzothiadiazole (BTH)-induced immunity (Nakayama et al., 2013). Overexpression of *TaWRKY45* in wheat confers enhanced protection against powdery mildew, leaf rust, and Fusarium head blight independently of specific *R* genes, and is highly responsive to SA and BTH (Bahrini, Ogawa, et al., 2011a; Bahrini, Sugisawa, et al., 2011; Heckmann et al., 2024a; Hu et al., 2019a). Similarly, *WRKY70* is essential for early NHP- induced SAR establishment and ROS bursts in Arabidopsis (Foret et al., 2024), while its cereal orthologs mediate broad-spectrum resistance and induced systemic resistance (ISR) (Andersen et al., 2020; H. Li et al., 2020; J. Li et al., 2004; M. Li et al., 2022a). The robust systemic induction of these specific WRKYs, alongside others linked to SA-responsive defense programs and stomatal immunity like *TaWRKY64* and *TaWRKY146* (Ma et al., 2017; Shah et al., 2023a; Walker et al., 2024a; Yang et al., 2023a), establishes that NHP-Induced Resistance (NIR) in wheat is primarily orchestrated through WRKY-dominated transcriptional networks, overlapping mechanistically with both Acquired Resistance (AR) and BTH-Induced Resistance (BIR) paradigms.

### Wheat deploys a distinct Pip-NHP-SA metabolic synergy

Exogenous NHP establishes a systemic metabolic state consistent with SAR-like immunity in wheat, characterized by a cooperative Pip–NHP–SA axis that is well-conserved across plant orders (Bernsdorff et al., 2016; Zeier, 2021). Consistent with observations in grasses where basal Pip levels are inherently higher and pathogen challenge triggers strong Pip induction (Lenk et al., 2019; Návarová et al., 2013; Sato et al., 2024; Schnake et al., 2020), local NHP application in wheat led to a sustained, systemic accumulation of Pip and NHP at 24 and 48 hpi. This metabolic state concurrently triggered a significant early spike in free SA and its 30-fold more abundant storage form, SAG, throughout systemic foliar tissue at 24 hpi, mirroring the broad SA-dependent transcriptional reprogramming of immune pathways observed in our dataset.

However, the biosynthetic origin of this SA surge highlights a key evolutionary divergence between eudicot and monocot SAR models. While Arabidopsis heavily relies on the isochorismate synthase (ICS1) pathway for pathogen-induced SA synthesis (Huang et al., 2020; Sun et al., 2015; Wang et al., 2011), wheat appears to favor the phenylalanine ammonia-lyase (PAL) pathway (Lefevere et al., 2020; Peng et al., 2021). In our study, no NHP-inducible ICS genes were detected; instead, several PAL transcripts were highly up-regulated, including the systemic induction of PAL2 and PAL4 orthologs by NHP. In Arabidopsis, PAL2 and PAL4 are required for full pathogen resistance by regulating metabolic flux toward SA and lignin biosynthesis (Allwood et al., 1999; Huang et al., 2010). Recent studies have further resolved the downstream enzymatic steps connecting phenylalanine-derived metabolism to SA biosynthesis in plants, including a BEBT–BBH–BSE module that converts benzoyl-CoA to SA (Y. Liu et al., 2025; Y. Wang et al., 2025; B. Zhu et al., 2025). Consistent with this pathway, NHP up- regulated wheat homologues of all three BEBT–BBH–BSE components. Together with increased SA and SAG accumulation, this supports the involvement of phenylalanine-derived SA biosynthesis during NHP-induced resistance, consistent with broader findings for PAL-driven SA accumulation in more pathogen-resistant cereal cultivars (Duan et al., 2014; Wu et al., 2023; Zhu et al., 2021).

Despite the strong accumulation of SA following NHP treatment, JA levels were not significantly altered, indicating no detectable SA–JA antagonism commonly reported in eudicots (Glazebrook, 2005; Pieterse et al., 2012; Van Der Does et al., 2013). This aligns with observations in rice, where over half of SA-inducible defense genes are also induced by JA (Tamaoki et al., 2013), and in barley, where Pip treatment does not compromise resistance against necrotrophic pathogens (Lenk et al., 2019). Furthermore, late-stage biotrophic infections in wheat have been shown to elevate ABA without suppressing NHP or SA accumulation (Sato et al., 2024). This suggests that NHP modulates JA signaling via WRKY and TIFY transcription factors rather than directly suppressing JA biosynthesis, granting cereals a flexible regulatory framework to simultaneously manage diverse biotrophic and necrotrophic stresses.

Finally, our data suggests that canonical lipid-transfer SAR signals deployed in Arabidopsis may be dispensable or even negatively regulated during NHP-mediated immunity in wheat. In Arabidopsis, glycerol-3-phosphate (G3P) and azelaic acid (AzA) act as mobile SAR signals dependent on lipid transfer proteins (LTPs) like DIR1 and EARLI1 for systemic transport (Cecchini et al., 2015; Chanda et al., 2011; Jung et al., 2009; Lim et al., 2016; Vlot et al., 2020; Wang et al., 2014; Wittek et al., 2014; Yu et al., 2013). Conversely, we found that NHP treatment during *Pt* infection strongly suppressed G3P metabolism and multiple DIR1-like and EARLI1-like LTP genes. This suppression mirrors findings that transient silencing of the wheat DIR1 ortholog, *TaDIR1-2*, enhances resistance against stripe rust, increasing H_2_O_2_ and SA accumulation while up-regulating *PR* genes and down-regulating ROS scavengers (Ahmed et al., 2017). The negative regulation of G3P and DIR1-like transcripts by NHP further emphasizes that wheat utilizes a distinct, potentially streamlined, signal mobilization network during systemic immunity.

### Evolutionary divergence and redundancy in the wheat NHP biosynthetic machinery

While core SAR components respond to NHP treatment in wheat, their regulation reflects lineage-specific expansion and specialization, and despite partial functional conservation, high and inducible expression of *TaSARD4* may not be required for NHP-mediated resistance in wheat against *Pt*. Identifying the functional wheat orthologues of the Arabidopsis NHP- biosynthetic enzymes (ALD1, SARD4, FMO1) has proven challenging due to the extensive expansion and redundancy of the FMO gene families in cereals (Gaba et al., 2023; Sun & Bakkeren, 2024; Thodberg & Jakobsen Neilson, 2020). Although the NHP pathway is broadly conserved across land plants (Schnake et al., 2020; Vlot et al., 2020), the polyploid wheat genome contains dozens of *ALD*-like aminotransferases and over 160 *TaFMO*-like candidates (Zhang et al., 2021). Consistent with recent transcriptomic evaluations (Zhang et al., 2021), *TaSARD4*-like homeologues and *ALD1*-like aminotransferases exhibit relatively weak or delayed transcriptional induction during early rust and powdery mildew infections (Sato et al., 2024). This suggests that, unlike the canonical Arabidopsis SAR model, NHP-mediated immunity in wheat during early *Pt* infection does not strictly rely on an early, massive transcriptional surge of *ALD1* and *SARD4*, central to Arabidopsis SAR. Instead, pathway redundancy, alternative enzymatic inputs, and cultivar-specific regulation likely obscure the contribution of individual biosynthetic components at the transcriptional level.

Functional characterization of the *TaSARD4* gene supported this interpretation; transiently silencing all three *TaSARD4* (A/B/D) homeologues using VIGS in Thatcher wheat did not significantly alter *Pt* disease progression during early infection. However, heterologous expression of the *TaSARD4-B* gene homeologue in the Arabidopsis *sard4* mutant partially restored NHP-*O*Glc accumulation, demonstrating that *TaSARD4-B* retains biochemical catalytic capacity. The lack of an altered resistance phenotype following transient *TaSARD4* knockdown, despite the enzyme’s functional competence, suggests that residual basal activity, highly redundant enzyme sets, or distinct, as-yet-uncharacterized routes may drive Pip and NHP production in wheat, mirroring speculation that alternative Pip-biosynthesis routes exist in Arabidopsis and other eudicots (Ding et al., 2016; Hartmann et al., 2017).

The genetic architecture of *TaFMO* candidates exhibits even more extensive diversification, spanning three main phylogenetic clades with massive lineage-specific expansions. While Arabidopsis relies heavily on FMO1 (from the B clade) for NHP biosynthesis, our RNA-seq analysis identified a single *TaFMO* gene from the A2 sub-clade (*TraesTC4B01G422800*) that was strongly and systemically up-regulated by NHP at 24 hpi. Notably, this gene is linked to the *Qfhs.ifa-5A* locus conferring resistance to *F. graminearum* (Nussbaumer et al., 2015). In contrast, two Group B *TaFMOs* (sub-clade B2) identified as DEGs in our dataset lacked an essential FMO catalytic motif (non-canonical), suggesting divergent functions or possible pseudogenization. The strong systemic induction of an A2 sub-clade *TaFMO* at 24 hpi which strictly precedes the significant NHP accumulation observed in distal leaves at 48 hpi, raises the possibility that this homeologue, rather than the canonical B-clade FMO1-like orthologs reported in Arabidopsis (Hartmann et al., 2018; Mishina & Zeier, 2006) and a previously identified B-clade *TaFMO* during seedling infection assays (Zhang et al., 2021), contributes to systemic *de novo* NHP biosynthesis in wheat.

These findings reinforce that the wheat NHP-biosynthetic pathway fundamentally diverges from the canonical Arabidopsis schema. Extensive wheat FMO redundancy and cultivar-dependent expression patterns may mask the direct transcriptional activation of genes like *TaALD1* and *TaSARD4*, reflecting a significant evolutionary divergence in how wheat and likely other monocots construct and deploy the NHP biosynthetic machinery. Resolving these relationships will require future pan-genome comparisons across resistant and susceptible wheat germplasm.

### A working model for NHP-Induced Resistance (NIR)

To synthesize our findings, we propose a working model for NHP-Induced Resistance (NIR) in hexaploid wheat, which integrates multi-tiered defense signaling across both local and systemic tissues (Figure 9). Historically, induced disease resistance in cereals has been characterized through two partially overlapping pathways: Acquired Resistance (AR), which relies heavily on localized, NPR1- and PR-dependent responses (Colebrook et al., 2012; Gao et al., 2018; Li et al., 2022), and Benzothiadiazole (BTH)-Induced Resistance (BIR), a chemically induced resistance response governed by WRKY transcriptional networks and Wheat- Chemically-Induced (WCI) defense genes with minimal reliance on canonical NPR1–PR pathways (Görlach et al., 1996; Li et al., 2020). Our data indicate that the NIR regulatory landscape bridges these two paradigms.

**Figure 9.**
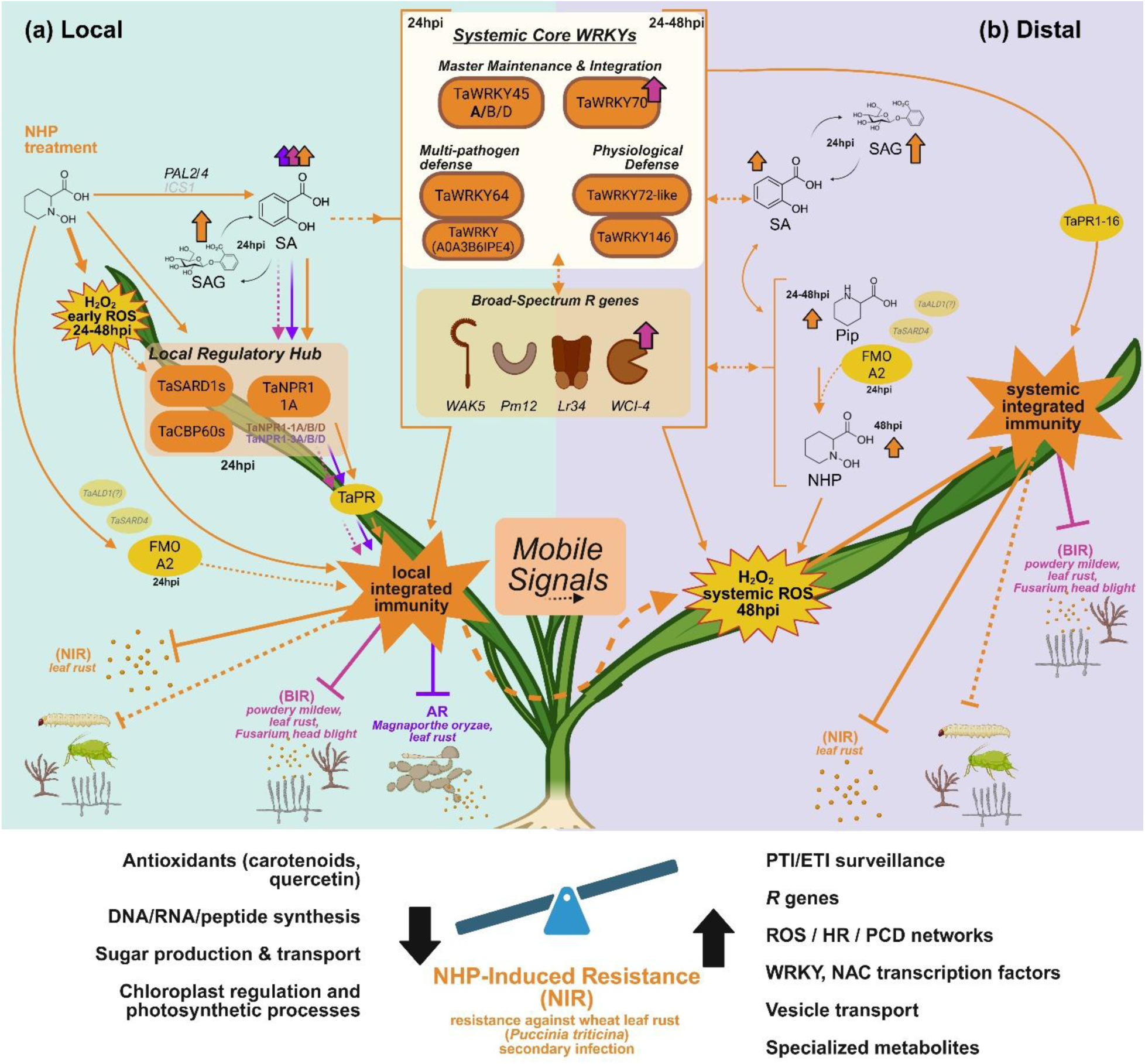
Proposed working model of NHP-Induced Resistance (NIR) and its integration with systemic immunity in hexaploid bread wheat. The NIR model (orange arrows) illustrates the spatio-temporal coordination of immune signaling following local NHP supplementation (10 mM) in the susceptible wheat cultivar Thatcher during early *Pt* pathogenesis. Pink arrows denote signaling components shared with the benzothiadiazole-induced resistance (BIR) pathway, while purple arrows highlight elements overlapping with acquired resistance (AR) described in cereals. Solid and dashed arrows correspond to strong, well-documented associations or inferred relationships, respectively. **(a) Local response (NHP-treated leaf)**. Local NHP treatment triggers whole-plant signaling characterized by an early ROS burst (24–48 hpi) and the activation of a local regulatory hub consisting of canonical SAR-regulatory elements (*TaSARD1s*, *TaCBP60s*, and *TaNPR1-1A*). NHP induces *PAL2* and *PAL4* (but not *ICS1*), driving high levels of SA and its 30-fold more abundant storage form (SAG), alongside the induction of a Group A *FMO-A2* from sub-clade A2 (without significant induction of *TaALD1*-like or *TaSARD4s*). The elevation in SA coincides with strong induction of systemic WRKY transcription factors and broad-spectrum resistance (*R*) genes in local leaves at 24 hpi, with *TaPR* genes likely promoted via the *TaNPR1-1A* hub, analogous to AR and BIR pathways. Together, these responses and the robust induction of PTI/ETI surveillance form the local integrated immunity that leads to enhanced restriction of *Pt*. Translocation of various mobile signals (which may include NHP, SA, and derivative compounds) moves systemically through the plant vasculature to prime distal leaves. **(b) Distal response (systemic priming in distal leaves)**. Systemic immunity is established by metabolic waves: an early SA/SAG surge (24 hpi) and early sustained Pip surge (24–48 hpi) are followed by a subsequent, slightly delayed NHP spike (48 hpi) correlating with a systemic ROS burst (48 hpi). These metabolic and ROS waves coordinate the systemic core WRKY hub (including major TFs *TaWRKY45* and *TaWRKY70*), which integrate SA/Pip/NHP signals to maintain defense and a heightened state of systemic immunity throughout 24–48 hpi. Concurrently, broad-spectrum *R* genes (*Lr34*, *Pm12*) and other relevant cereal SAR-associated defense genes (e.g., *WAK5*, *WCI-4*) contribute to the systemic restriction of *Pt* infection, interacting with the systemic core WRKYs and *PR* genes (e.g., *PR1- 16*) to initiate and maintain systemic integrated immunity. The bottom panel summarizes the reallocation of resources from primary growth toward enhanced PTI/ETI surveillance and whole- plant broad-spectrum immunity. **Alt text:** Two-panel comprehensive schematic model detailing spatial and temporal coordination of local defense hubs and distal priming during NHP-induced resistance in wheat.

Locally, NHP triggers an early AR-like response by prominently inducing a previously unreported *NPR1*-like homeologous triad alongside key SAR-regulatory elements such as *TaSARD1* and *TaCBP60s*, driving canonical downstream pathogenesis-related transcripts including *PR1-16* and *PR3* chitinases (Zhao et al., 2024). Concurrently, NIR initiates a BIR-like systemic signaling cascade by exclusively inducing over 50 WRKY transcription factors and the *WCI-4* gene (Zhang et al., 2021). This NIR state was characterized by the concurrent accumulation of SA, Pip, and NHP and strong systemic induction of WRKY transcription factors, most notably *TaWRKY45* and *TaWRKY70*, which have established roles in multi- pathogen defense (Bahrini, Ogawa, et al., 2011; Foret et al., 2024; Li et al., 2022).

Ultimately, NHP does not merely activate a single, specialized defense route. Instead, it functions as a master amplifier that reallocates primary metabolic resources toward heightened PTI/ETI surveillance, coordinated ROS waves, and broad-spectrum resistance networks. Understanding how the NIR pathway intersects with durable resistance (*R*) genes, such as the ABC transporter *Lr34* and the NLR *Pm12*, provides a compelling new avenue for agricultural biotechnology. By functionally characterizing and targeting these candidate regulatory components, including NHP-responsive *TaWRKYs* and cereal-specific *FMOs*, we can better leverage this endogenous systemic pathway. Collectively, our findings establish NHP as a master amplifier of systemic immunity in wheat, providing a mechanistic foundation for engineering durable, broad-spectrum disease resistance in global cereal crops.

## Supporting information

Supplemental Data Sets 1-5

## Supplementary Materials

### Supplemental Methods Documentation and Files

**Supplemental_Table_S1.docx** – Supplemental Table S1. Illumina paired-end RNA-seq library sizes, quality control parameters, and Kallisto pseudoalignment statistics.

**Supplemental_Data_Set_1.xlsx** – Supplemental Data Set 1. Experimental metadata and global transcriptomic differential gene expression profiles across basal infection and NHP-treatment conditions.

**Supplementary_Data_Set_2.xlsx** – Supplemental Data Set 2. Spatio-temporal functional co- expression subsets and Venn diagram intersections partition based on NHP-exogenous treatment and pathogen pressure.

**Supplemental_Data_Set_3.xlsx** – Supplemental Data Set 3. Gene Set Enrichment Analysis (GSEA) detailing enriched biological processes, normalized enrichment scores (NES), and leading-edge gene networks under NHP exposure.

**Supplemental_Data_Set_4.xlsx** – Supplemental Data Set 4. Weighted Gene Co-Expression Network Analysis (WGCNA) topological architecture and node properties of the highly NHP- responsive Blue module.

**Supplemental_Data_Set_5.xlsx** – Supplemental Data Set 5. Comprehensive list of diagnostic oligonucleotide primers, transgenic stocks, viral vectors, and experimental reagents utilized in this study.

### S1. Pathogen Propagation and Storage

*Pt* race 1 BBBD urediniospores were suspended at 5 mg/mL in Bayol F mineral oil and applied to wheat plants via airbrush at 150-200 psi. Inoculated plants were dried for 2-3 hours to evaporate the oil, incubated in a 100% relative humidity chamber in the dark for 16 hours, and subsequently moved to standard growth conditions. Urediniospores were collected from 7 to 14 days after infection on aluminum-covered trays. For short-term storage (<2 months), spores were maintained at 4°C in open tubes within a sealed container containing silica gel. For long-term storage, fresh spores were packaged in vegetarian pill capsules, desiccated under vacuum for 48 hours in the dark, and stored at −80°C in sealed screw-cap tubes. Prior to use, cryopreserved spores were rehydrated at room temperature (21°C) for 2-3 hours and heat-shocked in a 40°C water bath for 6 minutes.

### S2. RNA-Seq Bioinformatic Pipeline and Parameters

Following sequencing on the Illumina NovaSeq X-10B platform, raw sequence quality was assessed with FastQC (Andrews, 2010) and MultiQC (Ewels et al., 2016). Adapters and low- quality reads were removed using fastp v0.23.2. Cleaned reads were pseudo-aligned against the *T. aestivum* cv. Thatcher reference transcriptome utilizing Kallisto v0.50.2 with 100 bootstrap replicates. Crucially, the reference transcript FASTA file utilized for Kallisto index construction was structurally identical to the sequence database used for the comparative Liftoff annotation mapping, ensuring robust coordinate and ID consistency between differential gene expression tables and structural genomic loci. Transcript abundances were imported into R via the tximport package (Soneson et al., 2016). For global differential expression (comparing across broad conditions), the DESeq2 threshold was set at an adjusted *P*-value (FDR) < 0.05 and absolute log2 fold-change |L2FC| ≥ 1.5. For highly stringent filtering of specific spatial-temporal subsets, thresholds were adjusted to |L2FC| ≥ 2.0 and FDR ≤ 0.01. Network construction via WGCNA v1.72 utilized variance-stabilized counts following outlier removal, employing a signed network with a soft-thresholding power determined by scale-free topology criteria, and a minimum module size of 30.

### S3. Construct Design and Sequence Verification

Wheat cDNA was synthesized from RNA extracted from *Pt*-infected Thatcher seedlings using the SuperScript IV VILO Master Mix (ThermoFisher). Optimal regions for off-target-free silencing of all three *TaSARD4* homeologs were determined utilizing the siFi-21 algorithm (Lück et al., 2019). A 401-bp silencing fragment (corresponding to nucleotides 75-475) was amplified and cloned into the BSMV-γ vector (pCa-γbLIC) via ligation-independent cloning (LIC) (Pacak et al., 2010; Yuan et al., 2011). Overexpression constructs utilized the pCAMBIA1305-3HA vector backbone driven by the CaMV 35S promoter and terminated by the rbcS-E9t sequence. Plasmids were verified via Sanger sequencing prior to transformation into *Agrobacterium tumefaciens* strains C58 (for BSMV-VIGS) or GV3101/EHA105 (for stable *Arabidopsis* transformation).

### S4. Targeted LC-MS Phytohormone Extraction and Gradients (Wheat)

Frozen wheat leaf tissue (∼100 mg) was extracted overnight (18 hours, 200 rpm, 4°C) in a buffer containing 80% acetonitrile, 5% formic acid, 15% water, and 1 mM butylhydroxytoluene (BHT). Phase separation was induced by adding 40 mg MgSO₄ and 10 mg NaCl, vortexing, and centrifuging at 10,000 × g for 10 min. The supernatant was further dehydrated using 150 mg sodium sulfate, dried under a nitrogen gas stream, and redissolved in 1 mL of 1 M formic acid. Solid-phase extraction (SPE) was performed using Discovery DSC-18 SPE Tubes (500 mg, 3 mL; Supelco) conditioned with 100% methanol and 1 M formic acid. Metabolites were eluted in 2 mL 80% methanol, dried under nitrogen, and resuspended in 200 µL of 20% methanol for injection. Chromatographic separation was achieved on a Waters ACQUITY BEH C18 1.7 μm (2.1 x 50 mm) column at 35°C with a 0.3 mL/min flow rate. The mobile phases consisted of Optima-grade water with 0.1% formic acid (A) and acetonitrile with 0.1% formic acid (B). The gradient program was: 0-1 min 95% A; 1-9 min 95-50% A (curve 6); 9-11 min 50-10% A (curve 7); 11-13.5 min 10% A; 13.5-13.6 min 10-95% A (curve 6); and 13.6-16 min 95% A. The QTOF mass spectrometer operated in centroid MSe resolution mode with sequential positive (3500 V) and negative (−2500 V) electrospray ionization (ESI). Source parameters included: source temperature 125°C, desolvation temperature 300°C, sampling cone 40, cone gas 50 L/h, desolvation gas 600 L/h, and nebulizer 6.5 Bar. Analysis and absolute quantification against internal standards were conducted via MassLynx software.

## Author Contributions and Acknowledgements

S.S. designed and performed the majority of the experiments, analyzed the data, and wrote the manuscript. S.F. performed the lift-off mapping of Thatcher gene IDs to Chinese Spring gene IDs and assisted with troubleshooting RNA-seq pre-processing and pseudoalignments. J.H. additionally assisted in VIGS infection procedures on wheat plants and sample collection, contributing to the data in Figure 6. L.T. designed and performed experiments, C.H. analyzed the Arabidopsis metabolomic samples, generating the data used for Figure 7. C.C. designed the experimental workflow for targeted wheat metabolomics in foliar tissue, extracted samples alongside J.H., and provided guidance for peak integration and downstream analysis resulting in Figure 8. I.F., and Y.Z., provided strategic guidance and laboratory resources. G.B. attracted funding, provided resources and supervised the project and provided extensive editorial support.

We sincerely thank Dr. Corina Vlot and Dr. Robin Cameron for insightful discussions regarding SAR signaling in barley and Arabidopsis, as well as Dr. George Haughn and Dr. Xin Li for their rigorous evaluation and critical feedback which strengthened this work. We also acknowledge the Oklahoma Medical Research Foundation (OMRF) for RNA-sequencing services.

## Conflict of Interest

The authors declare that the research was conducted in the absence of any commercial or financial relationships that could be construed as a potential conflict of interest.

© 2026 by His Majesty the King in Right of Canada, as represented by the Minister of Agriculture and Agri-Food Canada, for the contribution by CC, JH and GB. © 2026 The authors SS, SF, CH, LT, SK, IF and YZ for their contribution. Published by Oxford University Press on behalf of American Society of Plant Biologists. This is an Open Access article distributed under the terms of the Creative Commons Attribution License (https://creativecommons.org/licenses/by/4.0/), which permits unrestricted reuse, distribution, and reproduction in any medium, provided the original work is properly cited.

---

**Supplementary Figure 1.**
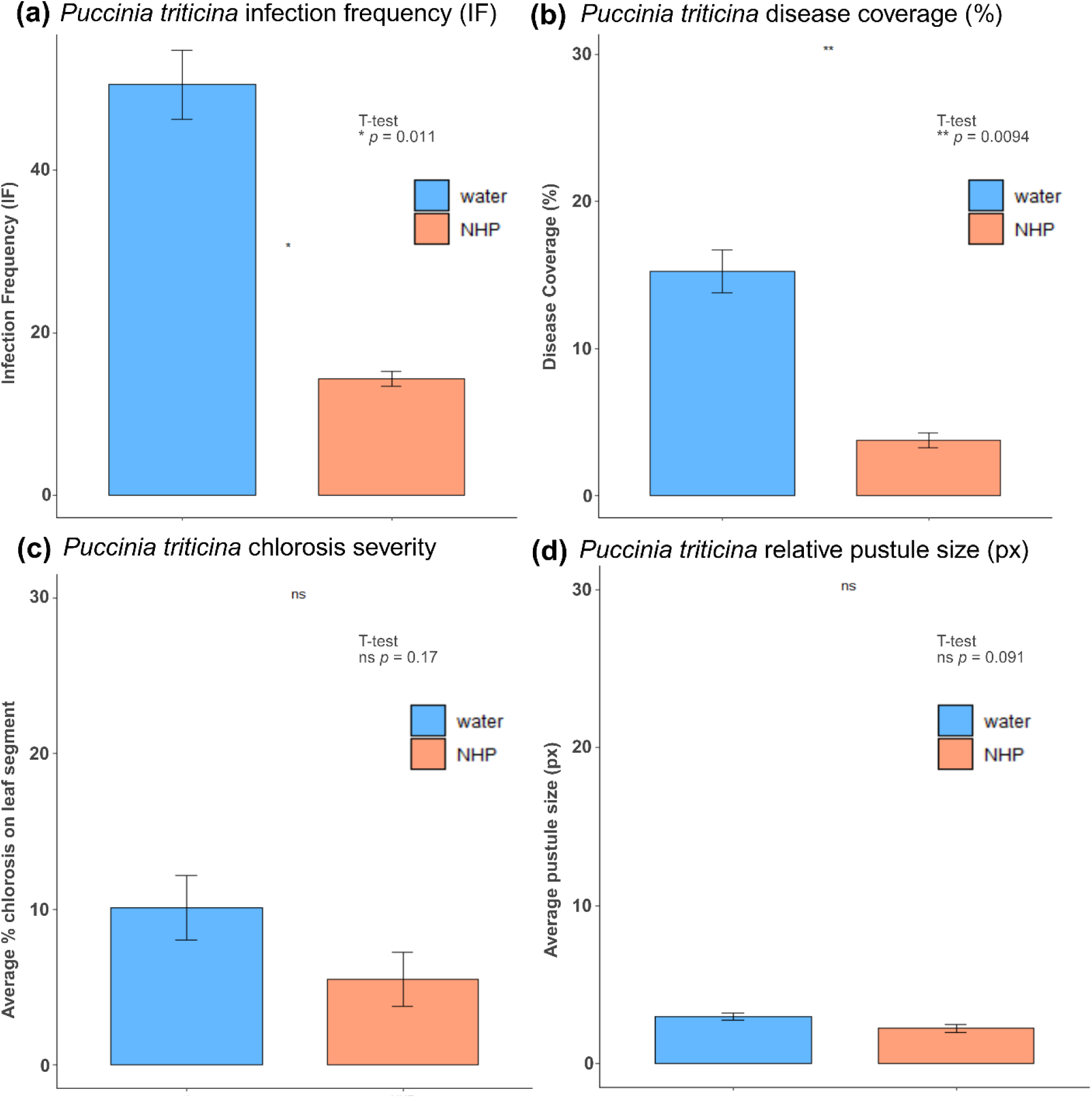
Effect of local *N*-hydroxypipecolic acid (NHP) pre-treatment on distal leaf disease parameters during *Pt* infection. Four-leaf stage susceptible wheat plants (cv. Thatcher) were locally infiltrated in the third leaf with a mock treatment (water) or 10 mM NHP. At 24 hours post-treatment, whole plants were systemically infected with *Pt* race 1 (BBBD) urediniospores. **(a–d)** Effect of local NHP treatment on distal (untreated) leaf disease parameters at 14 days after infection (DAI): **(a)** infection frequency (IF; pustules per cm^2^), **(b)** average % disease coverage, **(c)** average % chlorosis as measured by the semi-automated ImageJ macro designed to objectively quantify plant disease symptoms, Plant Immunity and Disease Image-based Quantification (PIDIQ), and **(d)** mean relative pustule size in pixels evaluated by the RUST algorithm. All bar plots represent mean values ± the standard error of the mean (SEM) from three independent experiments (*n* = 3 biological replicates), with each replicate comprising pooled samples from eight or nine individual plants. Asterisks indicate statistically significant differences determined by independent two-tailed Student’s t-tests: * *p* < 0.05, ** *p* < 0.01, ns = not significant.

**Supplementary Figure 2.**
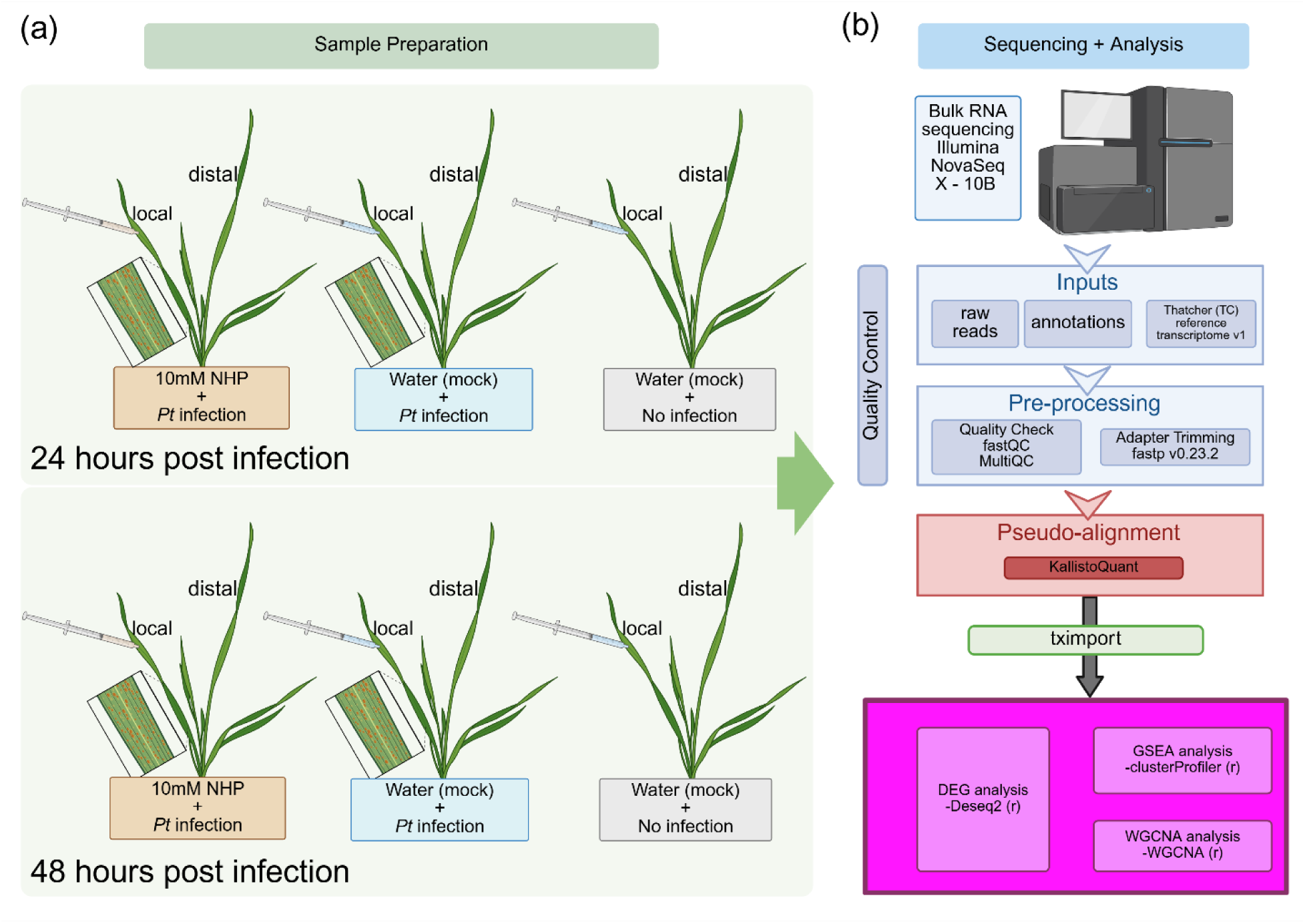
Experimental design and bioinformatic pipeline for global transcriptomic profiling of NHP-induced resistance. **(a)** Schematic representation of the experimental sample preparation. Four-leaf stage susceptible wheat plants (cv. Thatcher) were locally infiltrated with 10 mM NHP or a water (mock) control. At 24 hours post-treatment, plants were either systemically infected with *Pt* or left uninfected. Local treated leaves and distal untreated leaves were harvested at 24 and 48 hours post-infection (hpi). **(b)** RNA-sequencing and downstream bioinformatic analysis workflow. Bulk RNA was sequenced using the Illumina NovaSeq X-10B platform. Raw reads underwent quality control (FastQC, MultiQC) and adapter trimming (fastp v0.23.2). Cleaned reads were pseudo-aligned to the Thatcher reference transcriptome v1 using Kallisto. Transcript abundances were imported via tximport for downstream differential gene expression analysis (DESeq2), Gene Set Enrichment Analysis (clusterProfiler), and Weighted Gene Co-expression Network Analysis (WGCNA).

**Supplementary Figure 3.**
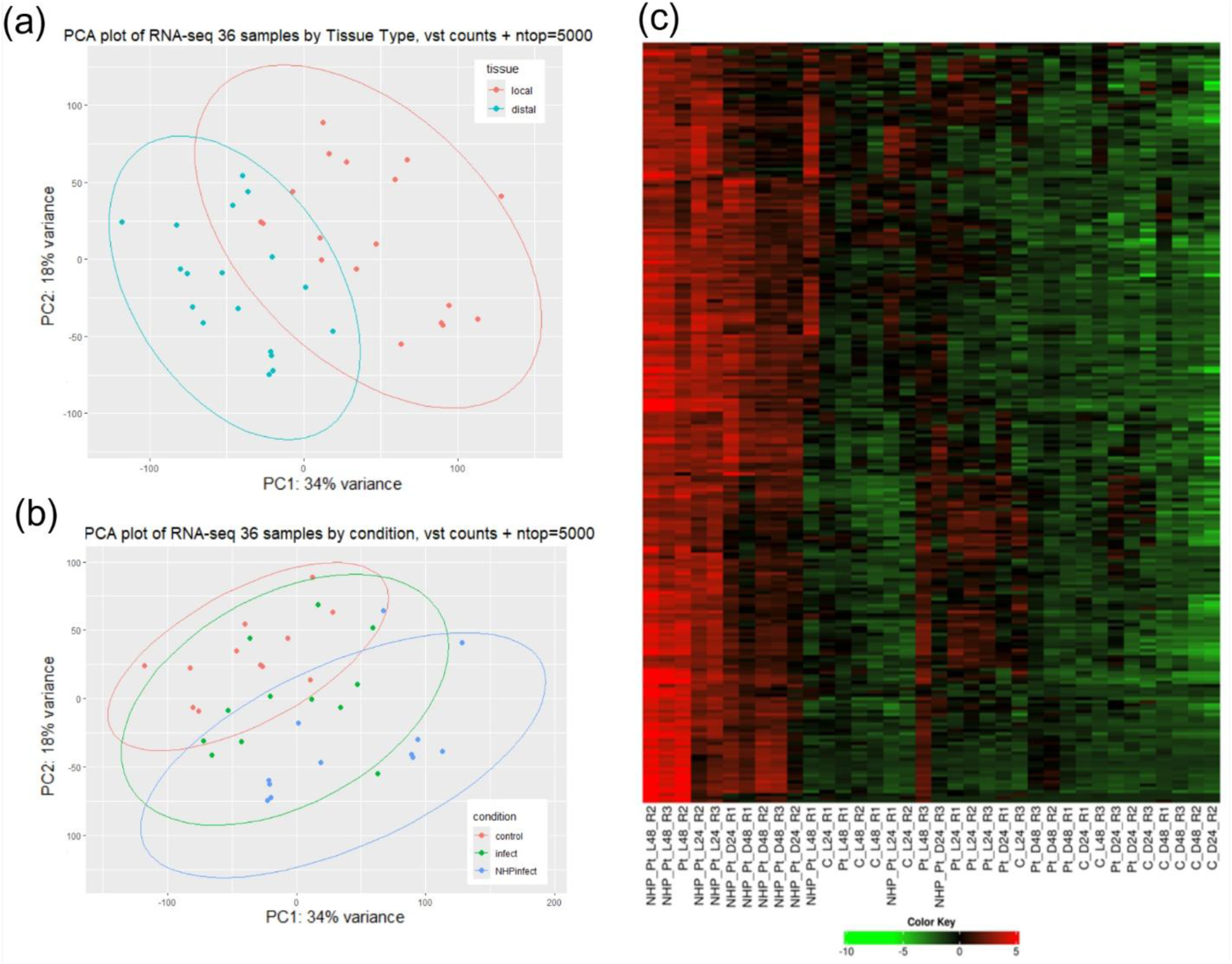
Exploratory analysis of RNA-seq data (36 samples). **(a)** Principal component analysis (PCA) biplot of the PC1 and PC2 axes for the top 5000 genes (ntop = 5000) subjected to variance-stabilized transformation (vst). Clear separation of tissue type (red for local leaves, blue for distal leaves) correlates with the PC1 axis, which accounts for 34% of the variation across samples. **(b)** PCA biplot of the PC1 and PC2 axes color-coded by treatment condition: control (uninfected, red), mock+Pt (water-treated, infected, green), and NHP+Pt (NHP-treated, infected, blue). **(c)** Unsupervised bi-cluster heatmap of all 36 RNA-seq samples, revealing the clustering of NHP-treated samples towards the left-most side of the plot (red color), and other samples associated with normal infection (mock+Pt) and control uninfected plants mostly clustering together (green color).

**Supplementary Figure 4.**
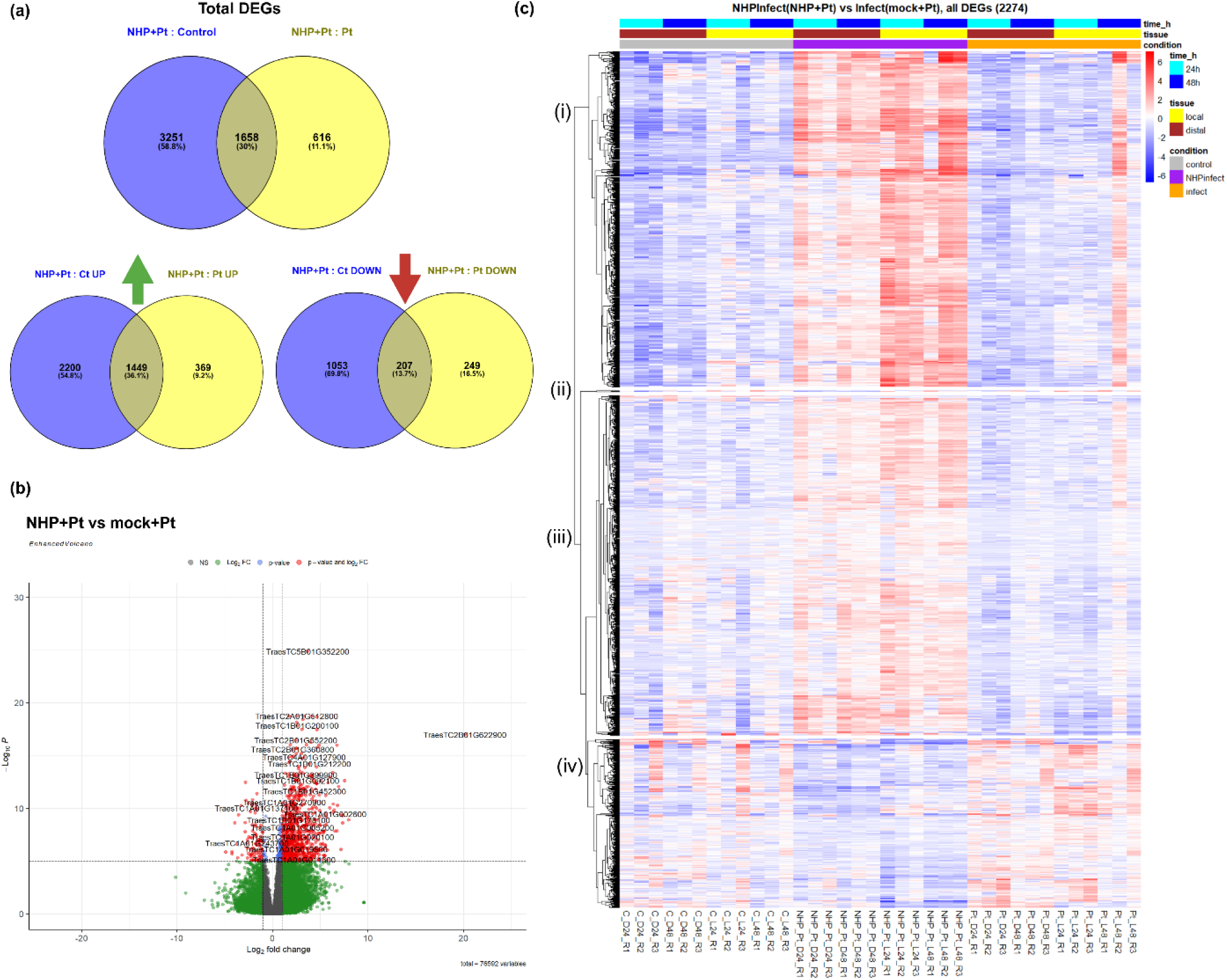
NHP remodels the global transcriptome during early *Pt* pathogenesis. **(a)** Venn diagram illustrating the overlap of total differentially expressed genes (DEGs) between NHP+*Pt* vs. control (blue) and NHP+*Pt* vs. mock+*Pt* (yellow) conditions. The 616 genes exclusive to the yellow circle represent DEGs specifically regulated by NHP supplementation during infection, but not significantly altered from an uninfected baseline. **(b)** Volcano plot of all 76,592 expressed genes comparing NHP+*Pt* to mock+*Pt* across all tissues and time-points. The x-axis represents Log₂ fold-change (L2FC) and the y-axis represents the - Log₁₀ adjusted *p*-value. Red dots denote significantly differentially expressed genes of interest (|L2FC| ≥ 1.5 and *p*adj < 0.05). **(c)** Hierarchical clustering heatmap of the 2,274 DEGs identified between NHP+*Pt* and mock+*Pt* conditions. Rows represent genes clustered based on Euclidean distances. Columns represent sample conditions: control uninfected (grey), NHP+*Pt* (purple), and mock+*Pt* (orange). Tissue types are indicated by yellow (local) and brown (distal) bars, and time-points are indicated by light blue (24 hpi) and dark blue (48 hpi) bars. Gene expression is displayed as a *z*-score from lower relative expression (dark blue) to higher relative expression (dark red).

**Supplementary Figure 5.**
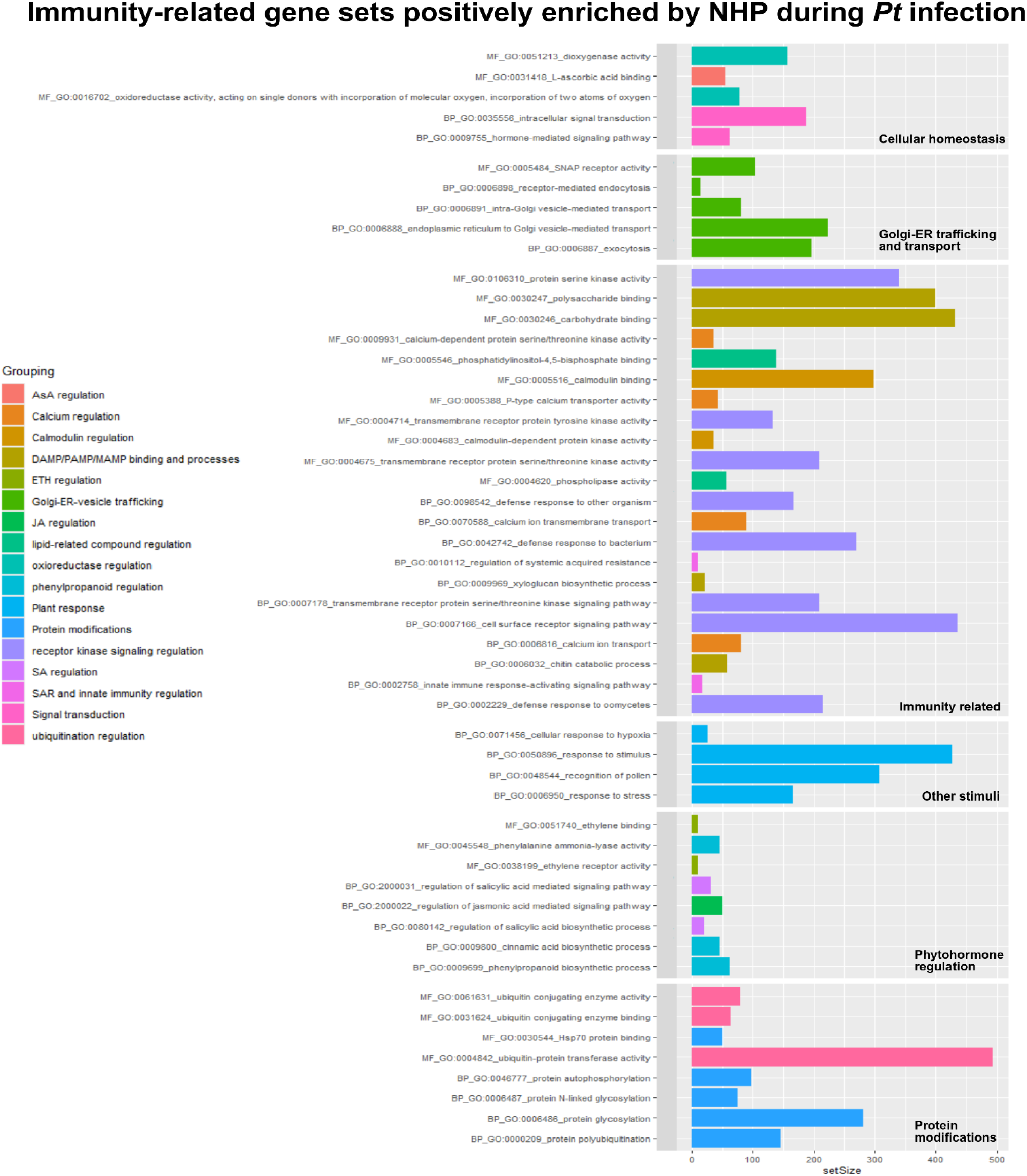
Immunity-related gene sets positively enriched upon NHP treatment. Gene sets shown represent significantly enriched (*p*adj < 0.05) categories derived from GSEA of all expressed genes comparing NHP+*Pt* vs mock+*Pt* conditions. Bar plot of significant gene sets categorized from top to bottom: cellular homeostasis, Golgi-ER-vesicle trafficking, immunity-related (PTI and ETI defense-related processes), other stimuli (abiotic), phytohormone regulation, and protein modifications. The *x*-axis refers to the number of genes reported in each gene set, and the color of each gene set is organized by broad functional annotations.

**Supplementary Figure 6.**
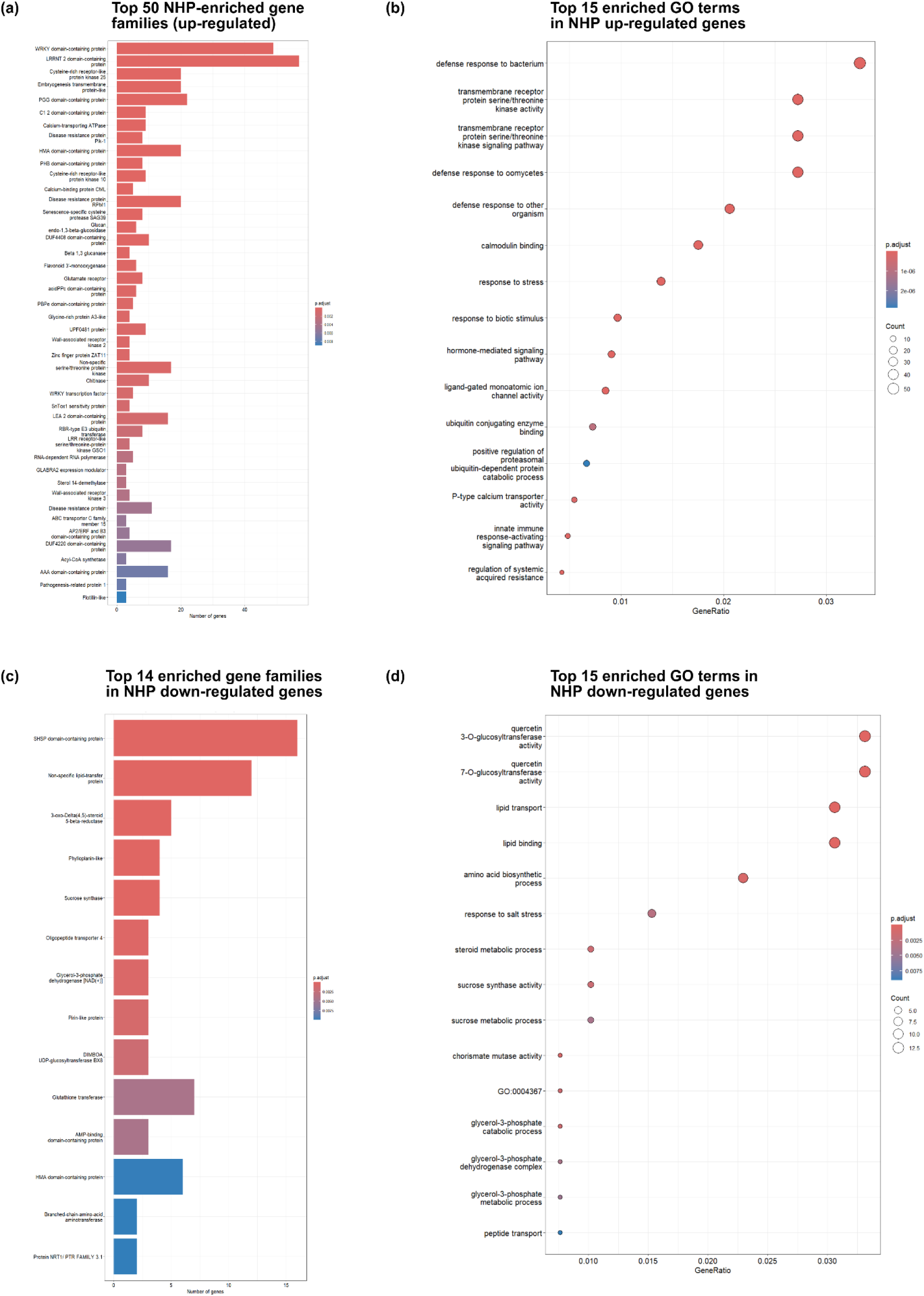
NHP systemically promotes defense-related gene families and represses primary cellular metabolism during *Pt* infection. Significantly enriched gene families and Gene Ontology (GO) terms derived from all expressed genes comparing NHP+*Pt* vs mock+*Pt* conditions. **(a)** Top 50 most significant gene families up-regulated in NHP-treated and *Pt*-infected leaves compared to a baseline *Pt* infection. **(b)** Top 15 most significantly positively enriched (up-regulated) GO terms. **(c)** Top 14 most significant gene families down-regulated in NHP+*Pt* conditions compared to a baseline *Pt* infection. **(d)** Top 15 most significantly negatively enriched (down-regulated) GO terms, highlighting the suppression of general primary metabolism. For bar plots (a, c), the x-axis displays the number of genes in each category. For bubble plots (b, d), the x-axis represents the gene ratio, and bubble size denotes the number of representative genes per category. Across all panels, color represents the adjusted p-value (< 0.05), with significance increasing as colors transition from blue to pink/red.

**Supplementary Figure 7.**
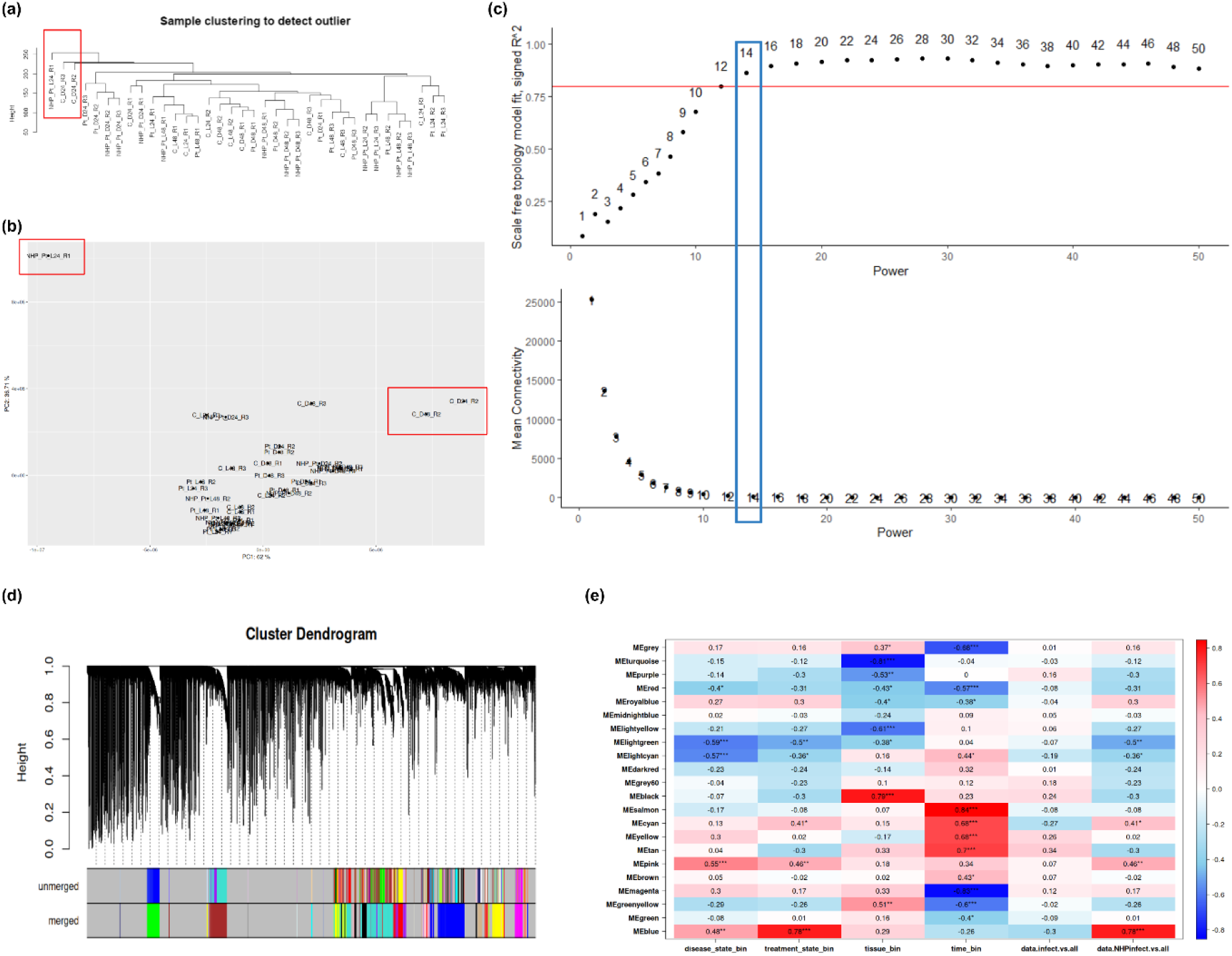
Whole-genome co-expression network analysis (WGCNA) module-trait correlation during early *Pt* pathogenesis. **(a)** Tree hierarchical clustering dendrogram of all RNA-seq samples used to identify potential outliers (highlighted in red). **(b)** PCA bi-cluster plot confirming separation of identified outlier samples (red boxes). **(c)** Soft- threshold power selection based on scale-free topology criterion (signed R2) and mean connectivity; a power of 14 (blue box) was selected for network construction. **(d)** Gene dendrogram showing unmerged (top) and merged (bottom) co-expression modules used for downstream analyses. **(e)** Module–trait correlation heatmap for 22 co-expression modules, showing correlations between module eigengenes and experimental variables (disease state, treatment, tissue type, timepoint). Correlation coefficients are displayed (−0.8 to 0.8), with color scale indicating strength and direction of association. The MEblue module, showing strong positive association with NHP treatment, was selected for further network and hub-gene analysis.

**Supplementary Figure 8.**
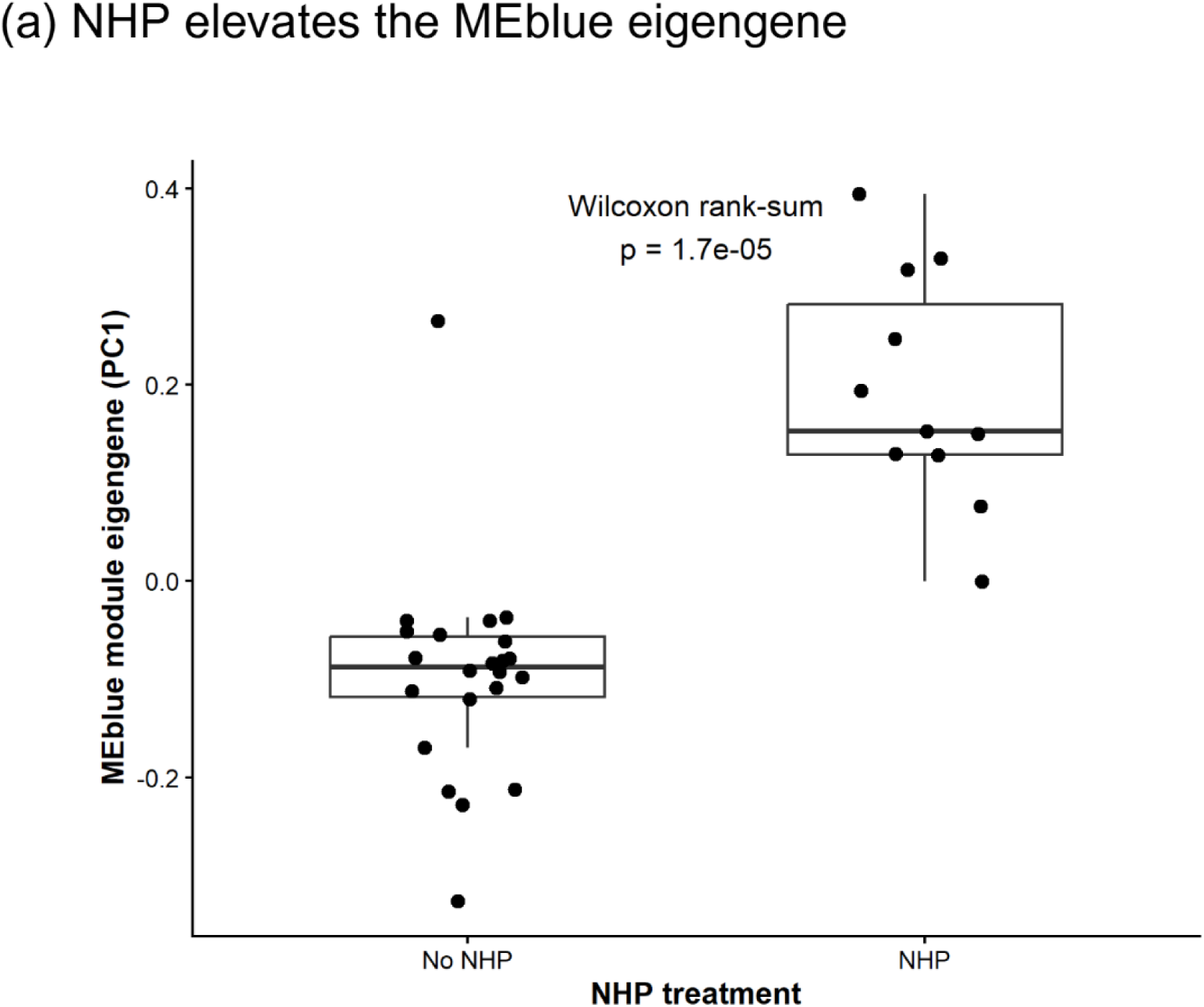
NHP-associated co-expression network of the MEblue module. (a) MEblue module eigengene values (PC1) across samples with and without NHP treatment. MEblue eigengene expression was significantly elevated in NHP-treated samples (Wilcoxon rank-sum test, *p* = 1.7 × 10⁻^5^), indicating coordinated transcriptional activation associated with NHP exposure.

**Supplementary Figure 9.**
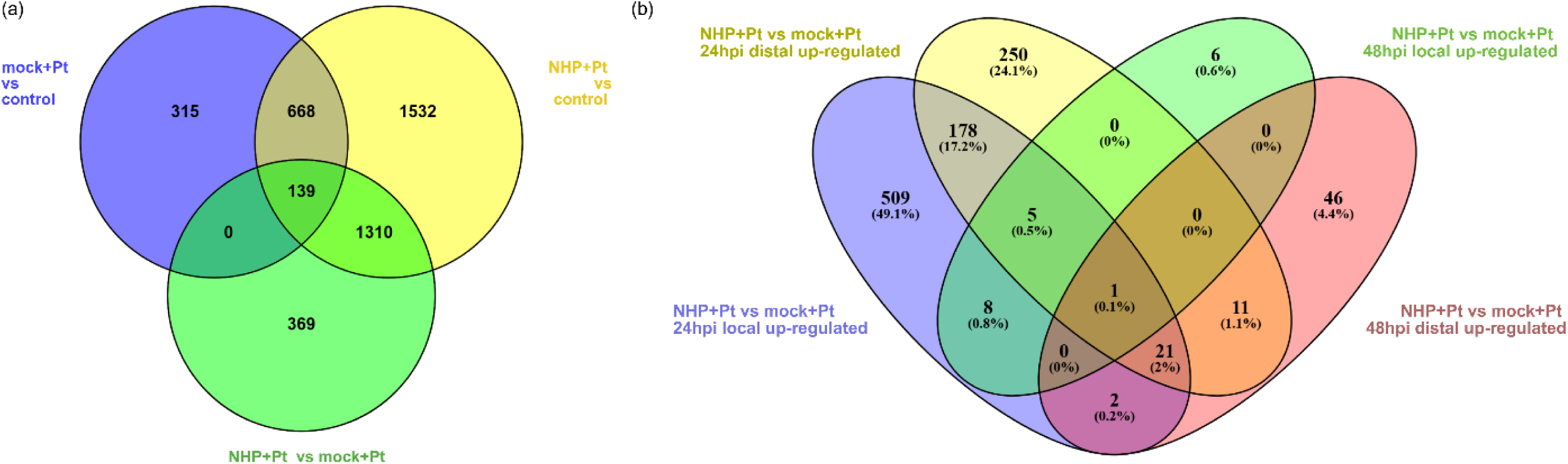
Spatial and temporal overlap of NHP-induced differentially expressed genes (DEGs). **(a)** Venn diagram illustrating the overlap of significantly up-regulated DEGs across three pair-wise comparisons: mock+*Pt* vs. control (blue), NHP+*Pt* vs. control (yellow), and NHP+*Pt* vs. mock+*Pt* (green). The overlap between the yellow and green circles represents genes specifically induced by NHP during *Pt* infection. **(b)** Venn diagram detailing the temporal (24 and 48 hpi) and spatial (local treated and distal untreated leaves) intersections of the up-regulated DEGs specifically from the NHP+*Pt* vs. mock+*Pt* comparison, delineating genes commonly up-regulated to establish the systemic NHP footprint.

**Supplementary Figure 10.**
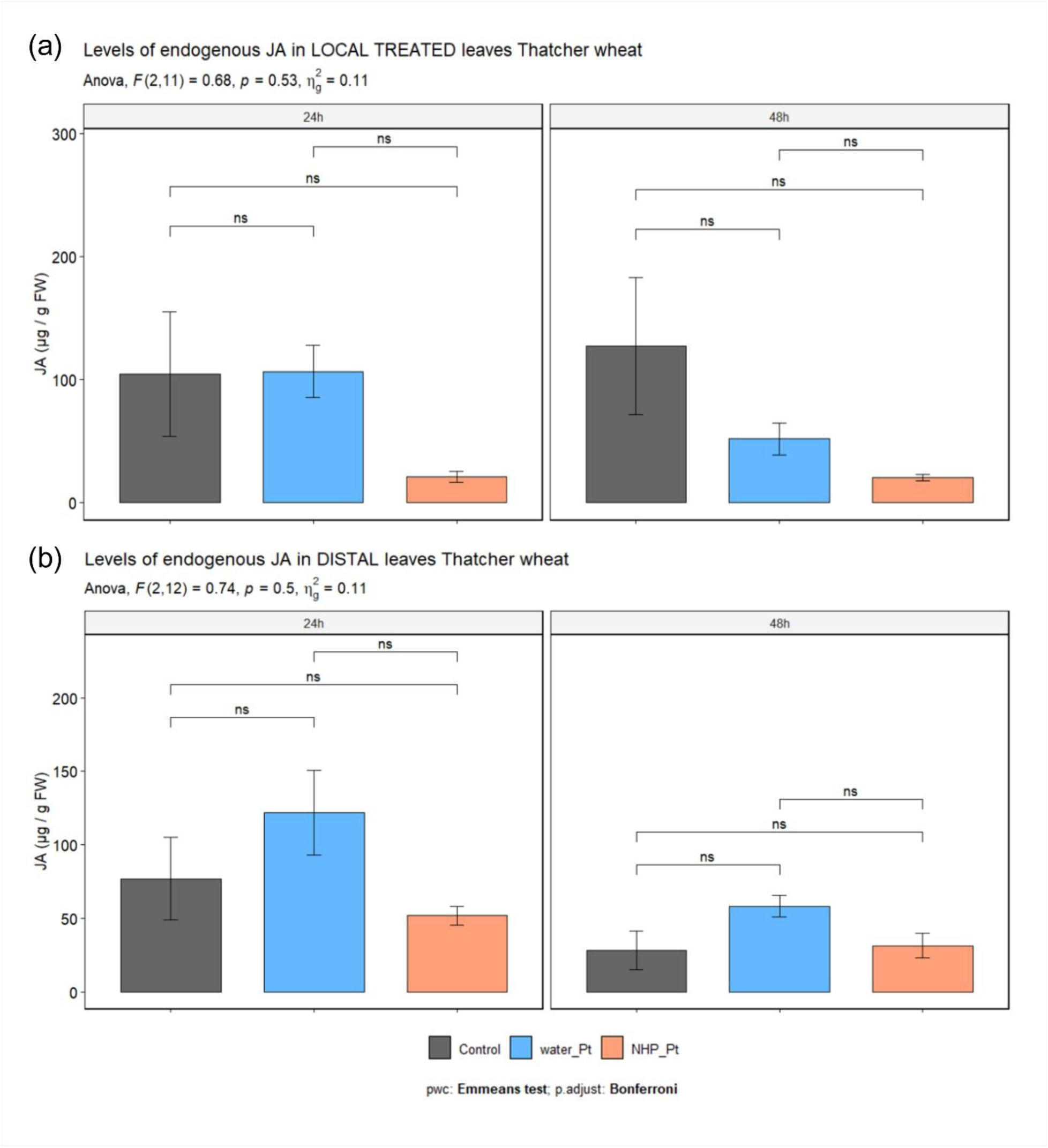
Local NHP supplementation does not significantly impact JA concentrations across treatments and time points. Four-leaf stage Thatcher wheat plants were infiltrated in the second true leaf with either a mock treatment (water) or 10 mM NHP. At 24 hours post-treatment, plants were either systemically inoculated with *Pt* race 1 BBBD urediniospores or left uninfected (control). Local treated leaves and distal untreated leaves were harvested at 24 and 48 hours post-infection (hpi) for targeted LC-MS analysis. Bar plots display concentrations (μg/g fresh weight) for local treated leaves (**a,** top row) and distal untreated leaves (**b,** bottom row) at 24 and 48 hpi. Grey bars represent uninfected controls, blue bars represent mock-treated infected plants, and orange bars represent NHP-treated infected plants. Graph data are presented as mean values ± SEM from three independent experiments (*n* = 3 biological replicates), with each replicate pooling tissue from five individual plants. A two-way ANOVA (Type II test) with Bonferroni post-hoc pairwise comparisons were conducted (* *p* < 0.05, ** *p* < 0.01, *** *p* < 0.001, **** *p* < 0.0001; *nd* = not detected; blank = not significant.

**Supplementary Figure 11.**
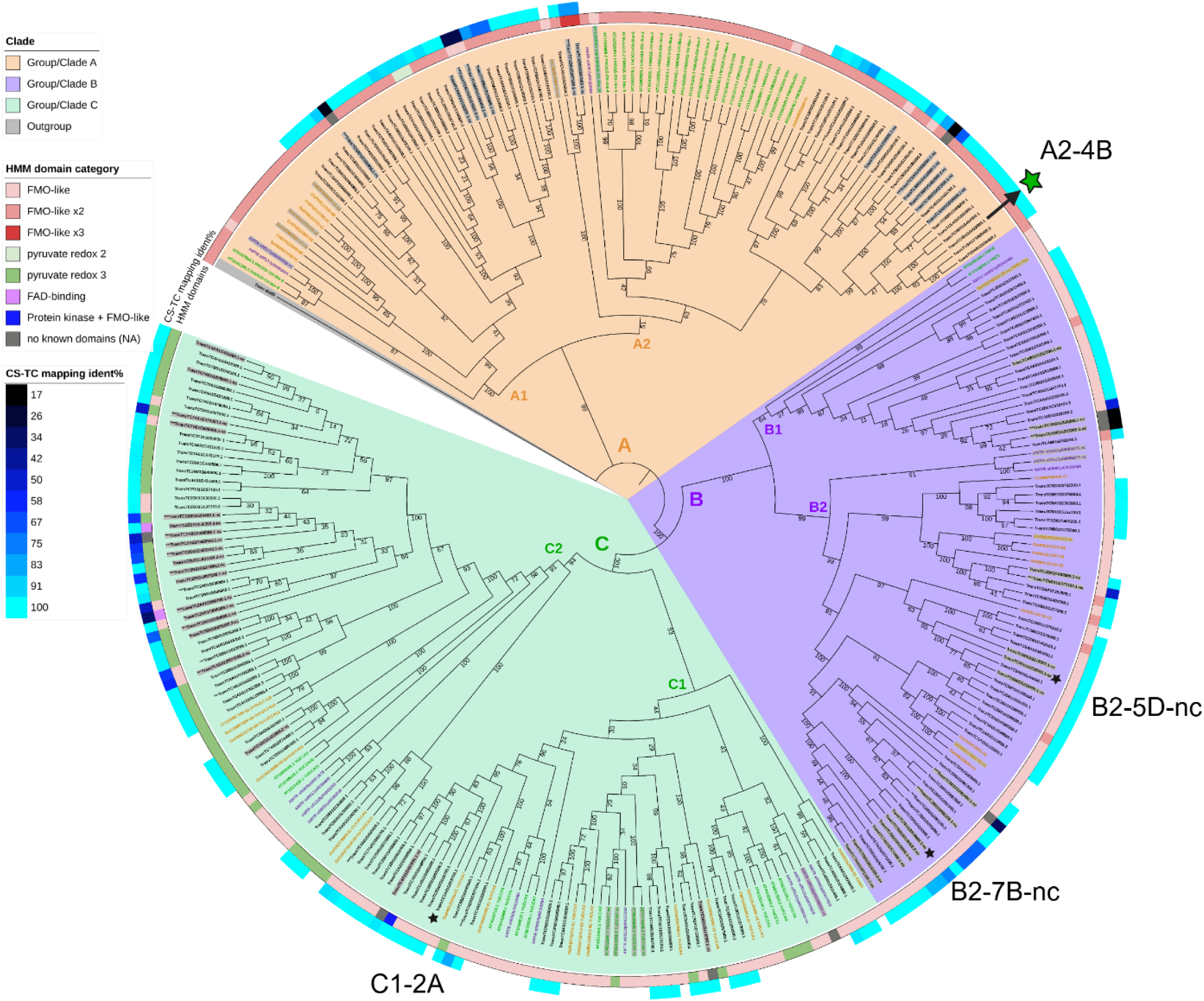
Phylogenetic distribution and NHP-responsiveness of the highly expanded Flavin-Dependent Monooxygenase (FMO) family in wheat cv. Thatcher. Maximum-likelihood phylogenetic tree aligning 163 *TaTC-FMO* proteins from wheat cv. Thatcher with orthologs from *Arabidopsis thaliana* (green text), *Amborella trichopoda* (purple text), and *Oryza sativa* ssp. *japonica*, utilizing *Galdieria sulphuraria* as an outgroup (grey clade). The inner tree denotes the major TaFMO clades A, B, and C. The inner colored track signifies predicted hidden Markov model (HMM) domain categories, while the outer blue gradient track indicates the percentage of protein sequence identity between Thatcher FMOs and Chinese Spring reference FMOs. Gene IDs highlighted in grey lack conserved canonical FMO motifs. Star symbols denote differentially expressed *TaTC-FMOs* detected via RNA-seq during early *Pt* pathogenesis; the green star specifically highlights *TraesTC4B01G422800* (sub-clade A2), which was systemically up-regulated by local NHP pre-treatment.

**Supplementary Figure 12.**
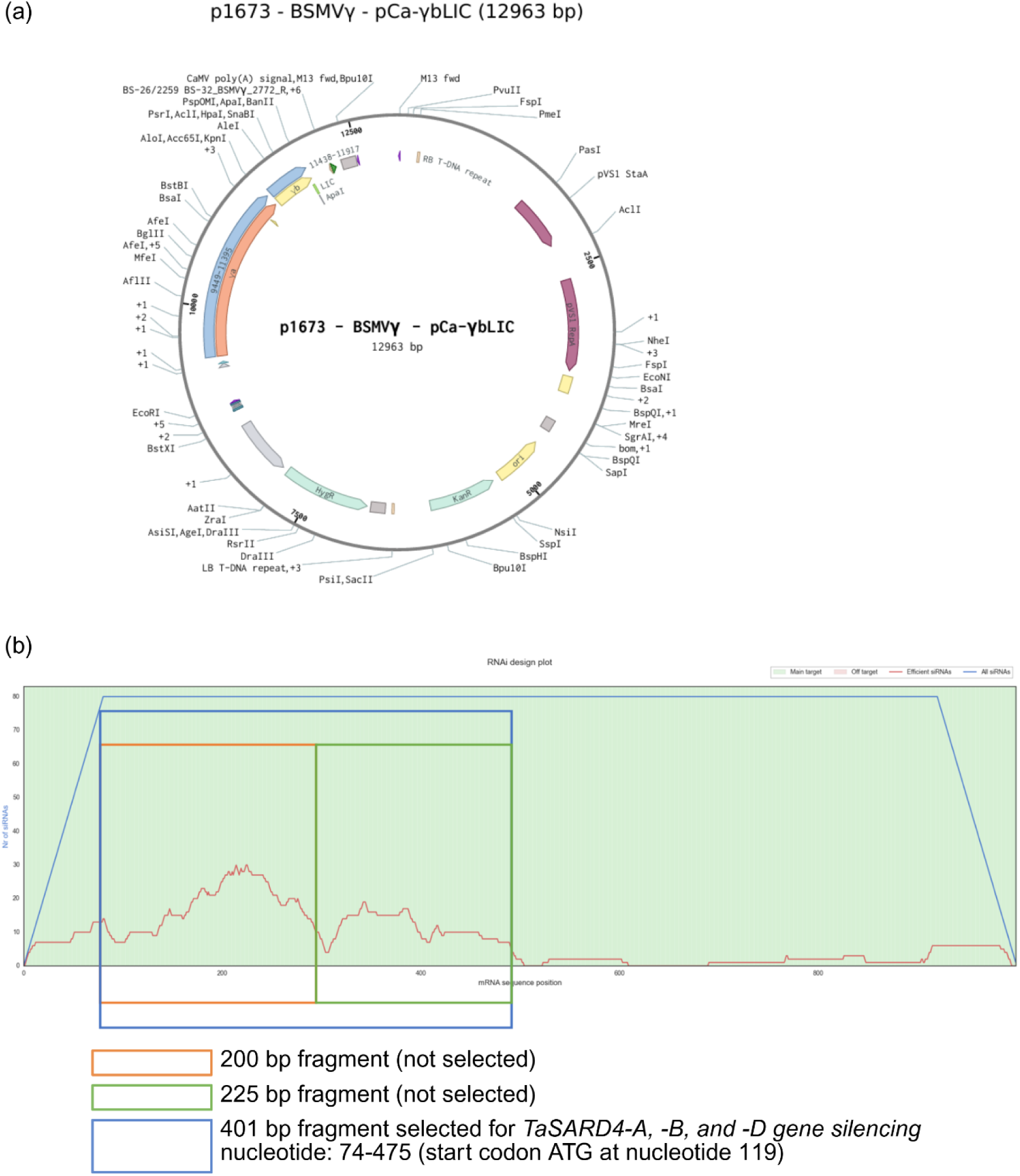
Virus-induced gene silencing (VIGS) fragment design and construct. **(a)** A plasmid map of the BSMV- *γ* plasmid (p2300, derived from a p1673 backbone) used for ligation independent cloning (LIC) of *TaSARD4* Thatcher fragment (401bp) insertion at the LIC site for transient delivery via *Agrobacterium tumefaciens* strain C58 into tobacco for viral propagation. **(b)** Fragment construct target area of a 401bp fragment (blue box) spanning the 5’UTR region, ATG start codon, and beginning half of the *TaSARD4* gene; designed using the siFi-21 algorithm for use in silencing. Red line indicates efficiency of small interfering RNA (siRNA) regions for all three Thatcher *TaSARD4* target sequences (*TaSARD4-A*, -*B*, and -*D*) against protein-encoding genes in Thatcher background. Sequence verified via in-house sanger sequencing: TCGCGAACGCCGCGAGCAGAGCGGGAGCAGCCCCACCGCGGCAGCATGGCAGCGC CGCCGTCCCACCCGTTCGTCTACATCGACGCCGAGGCGCTCCACGCCGTCCTCCCCT TCCCGTCCCTGATCTCCCACCTCCGCGCGGGCCTCCCCGCCTTCTCCGGTGACATCCA CTGCCCCCACCGCGTCTCCTTCCCGGTCCCGACCTCGCCCTCCGCGGCGTTGCTCCTC ATGCCCTCATGGTGCGCCCACCCCTCGCTCCCCTACCTCGCCCTCAAGGCCGTCACC TCCTTCCCCGCCAACTCCCCGCACCGCCCCTCCATCCACGCCACCGTCTCCCTCTTCA GCTCCGCCTCCGGCGTGCCCCTCGCGTCCGTCGACGGCTCCGCGCTCACCCTCCTCCG

