## Supplementary Table S1. for "*N*-Hydroxypipecolic Acid Triggers Systemic Acquired Resistance Against Leaf Rust (*Puccinia triticina*) in Wheat"

Supplementary Table S1. Library sizes of 36 RNA-seq samples Illumina (paired-end): QC and Kallisto pseudoalignment stats (bioproject accession: PRJNA1285943, study: SRP598210)

| library_ID | | | | | | | | | | | Kallisto pseudoalignment QC | | |
|---|---|---|---|---|---|---|---|---|---|---|---|---|---|
| Sample_Name | library_ID | accession | biosample_accession | raw_reads | raw_bases_(Gb) | clean_reads | clean_bases_(Gb) | Q20_% | Q30_% | GC_% | n_processed | n_pseudoaligned | %_pseudoaligned |
| C_L24_R1 | C-L1_24h | SRR34364317 | SAMN49784114 | 13618695 | 20.56423 | 1.33E+08 | 17.41089 | 97.19026 | 92.20814 | 56.30621 | 66528814 | 54568707 | 82.0 |
| C_L24_R2 | C-L2_24h | SRR34364316 | SAMN49784115 | 1.47E+08 | 22.18509 | 1.44E+08 | 19.10986 | 97.02134 | 91.81646 | 56.19522 | 71854995 | 58687756 | 81.7 |
| C_L24_R3 | C-L3_24h | SRR34364305 | SAMN49784116 | 1.4E+08 | 21.15184 | 1.37E+08 | 18.2972 | 95.90373 | 89.48041 | 58.07924 | 68453807 | 54547446 | 79.7 |
| C_D24_R1 | C-D1_24h | SRR34364294 | SAMN49784117 | 45395498 | 6.85472 | 44463932 | 5.849346 | 97.53529 | 93.13143 | 55.26769 | 22697749 | 17955298 | 79.1 |
| C_D24_R2 | C-D2_24h | SRR34364287 | SAMN49784118 | 1.57E+08 | 23.67434 | 1.53E+08 | 20.53657 | 97.44475 | 92.73249 | 54.51569 | 76690141 | 64753104 | 84.4 |
| C_D24_R3 | C-D3_24h | SRR34364286 | SAMN49784119 | 99002678 | 14.9494 | 96846220 | 12.57142 | 97.6108 | 93.2476 | 54.90695 | 48423110 | 41318200 | 85.3 |
| Pt_L24_R1 | Mock-Pt-L1_24h | SRR34364285 | SAMN49784120 | 1.73E+08 | 26.19181 | 1.7E+08 | 22.12913 | 97.65645 | 93.3263 | 55.42559 | 84784578 | 72740265 | 85.8 |
| Pt_L24_R2 | Mock-Pt-L2_24h | SRR34364284 | SAMN49784121 | 1.27E+08 | 19.15037 | 1.24E+08 | 17.06797 | 97.89444 | 93.9143 | 54.74447 | 63441634 | 53566303 | 84.4 |
| Pt_L24_R3 | Mock-Pt-L3_24h | SRR34364283 | SAMN49784122 | 1.98E+08 | 29.96026 | 1.95E+08 | 27.69872 | 93.04341 | 83.03561 | 66.79322 | 97353172 | 36537414 | 37.5 |
| Pt_D24_R1 | Mock-Pt-D1_24h | SRR34364282 | SAMN49784123 | 1.87E+08 | 28.18785 | 1.83E+08 | 24.615 | 97.59178 | 93.08275 | 54.89771 | 91437020 | 78755804 | 86.1 |
| Pt_D24_R2 | Mock-Pt-D2_24h | SRR34364315 | SAMN49784124 | 1.57E+08 | 23.72356 | 1.54E+08 | 21.6059 | 93.21582 | 83.64746 | 64.27441 | 76869917 | 36070913 | 46.9 |
| Pt_D24_R3 | Mock-Pt-D3_24h | SRR34364314 | SAMN49784125 | 66910704 | 10.10352 | 65630220 | 8.829596 | 97.70185 | 93.3489 | 53.25496 | 33455352 | 26424605 | 79.0 |
| NHP_Pt_L24_R1 | NHP-Pt-L1_24h | SRR34364313 | SAMN49784126 | 81161612 | 12.25993 | 79575770 | 11.75043 | 86.78033 | 70.63361 | 70.14162 | 39787885 | 19443115 | 48.9 |
| NHP_Pt_L24_R2 | NHP-Pt-L2_24h | SRR34364312 | SAMN49784127 | 1.37E+08 | 20.74689 | 1.35E+08 | 18.05552 | 97.72564 | 93.42121 | 54.63641 | 67351155 | 58297746 | 86.6 |
| NHP_Pt_L24_R3 | NHP-Pt-L3_24h | SRR34364311 | SAMN49784128 | 1.77E+08 | 26.78347 | 1.73E+08 | 21.44232 | 97.79057 | 93.75251 | 55.50477 | 86585020 | 75422715 | 87.1 |
| NHP_Pt_D24_R1 | NHP-Pt-D1_24h | SRR34364310 | SAMN49784129 | 1.57E+08 | 23.64326 | 153.133 | 20.98104 | 97.45997 | 92.91796 | 54.0903 | 78288934 | 61506197 | 78.6 |
| NHP_Pt_D24_R2 | NHP-Pt-D2_24h | SRR34364309 | SAMN49784130 | 1.5E+08 | 22.59425 | 1.46E+08 | 20.72349 | 94.37134 | 86.04083 | 62.55788 | 73201599 | 37757476 | 51.6 |
| NHP_Pt_D24_R3 | NHP-Pt-D3_24h | SRR34364308 | SAMN49784131 | 1.69E+08 | 25.50155 | 1.66E+08 | 23.85406 | 91.63555 | 80.13342 | 68.1139 | 82906693 | 26068133 | 31.4 |
| C_L48_R1 | C-L1_48h | SRR34364307 | SAMN49784132 | 1.36E+08 | 20.54064 | 1.33E+08 | 17.5053 | 97.58449 | 93.22633 | 55.22683 | 66641474 | 57584617 | 86.4 |
| C_L48_R2 | C-L2_48h | SRR34364306 | SAMN49784133 | 1.03E+08 | 15.52402 | 1.01E+08 | 13.64847 | 97.54943 | 93.149 | 54.80161 | 50454492 | 43587437 | 86.4 |
| C_L48_R3 | C-L3_48h | SRR34364304 | SAMN49784134 | 1.44E+08 | 21.67077 | 1.41E+08 | 19.54275 | 95.45486 | 88.65448 | 59.03127 | 70356939 | 48908789 | 69.5 |
| C_D48_R1 | C-D1_48h | SRR34364303 | SAMN49784135 | 1.26E+08 | 19.00424 | 1.23E+08 | 17.30107 | 94.22511 | 85.74453 | 60.32489 | 61556899 | 36962164 | 60.0 |
| C_D48_R2 | C-D2_48h | SRR34364302 | SAMN49784136 | 1.33E+08 | 20.01881 | 1.3E+08 | 17.32686 | 97.51688 | 93.04795 | 54.18404 | 65088747 | 55652560 | 85.5 |
| C_D48_R3 | C-D3_48h | SRR34364301 | SAMN49784137 | 1.8E+08 | 27.1477 | 1.76E+08 | 24.78916 | 93.85104 | 84.91975 | 63.05862 | 87876786 | 47457002 | 54.0 |
| Pt_L48_R1 | Mock-Pt-L1_48h | SRR34364300 | SAMN49784138 | 1.65E+08 | 24.9498 | 1.62E+08 | 22.57771 | 96.22131 | 90.14802 | 58.49591 | 81112350 | 57974359 | 71.5 |
| Pt_L48_R2 | Mock-Pt-L2_48h | SRR34364299 | SAMN49784139 | 1.19E+08 | 17.98592 | 1.17E+08 | 15.92168 | 96.04088 | 89.86352 | 58.26513 | 58450144 | 44961556 | 76.9 |
| Pt_L48_R3 | Mock-Pt-L3_48h | SRR34364298 | SAMN49784140 | 75516430 | 11.40298 | 74131516 | 10.29691 | 97.84248 | 93.64661 | 54.57098 | 37065758 | 32704286 | 88.2 |
| Pt_D48_R1 | Mock-Pt-D1_48h | SRR34364297 | SAMN49784141 | 1.42E+08 | 21.45137 | 1.39E+08 | 19.87691 | 97.72867 | 93.36526 | 53.91512 | 69591714 | 59986831 | 86.2 |
| Pt_D48_R2 | Mock-Pt-D2_48h | SRR34364296 | SAMN49784142 | 1.02E+08 | 15.43364 | 1E+08 | 14.1838 | 94.40814 | 86.36911 | 60.68383 | 50023502 | 30316436 | 60.6 |

| | | | | | | | | | | | | | |
|---|---|---|---|---|---|---|---|---|---|---|---|---|---|
| Pt_D48_R3 | Mock-Pt-D3_48h | SRR34364295 | SAMN49784143 | 93121546 | 14.06135 | 91343714 | 12.35301 | 97.22456 | 92.43697 | 55.38483 | 45671857 | 38401780 | 84.1 |
| NHP_Pt_L48_R1 | NHP-Pt-L1_48h | SRR34364293 | SAMN49784144 | 1.7E+08 | 25.63309 | 1.67E+08 | 23.06795 | 97.82863 | 93.61235 | 53.66596 | 83271057 | 72566799 | 87.1 |
| NHP_Pt_L48_R2 | NHP-Pt-L2_48h | SRR34364292 | SAMN49784145 | 1.77E+08 | 26.67813 | 1.73E+08 | 24.11501 | 95.84118 | 89.3671 | 59.17707 | 86737174 | 61949909 | 71.4 |
| NHP_Pt_L48_R3 | NHP-Pt-L3_48h | SRR34364291 | SAMN49784146 | 1.43E+08 | 21.62463 | 1.4E+08 | 19.39634 | 97.08538 | 92.09014 | 56.22998 | 70244744 | 56997069 | 81.1 |
| NHP_Pt_D48_R1 | NHP-Pt-D1_48h | SRR34364290 | SAMN49784147 | 1.42E+08 | 21.43179 | 1.39E+08 | 19.30344 | 97.76794 | 93.46392 | 53.11367 | 69589279 | 60244604 | 86.6 |
| NHP_Pt_D48_R2 | NHP-Pt-D2_48h | SRR34364289 | SAMN49784148 | 82054684 | 12.39026 | 80506144 | 10.84187 | 97.32848 | 92.61987 | 56.01895 | 40253072 | 32901623 | 81.7 |
| NHP_Pt_D48_R3 | NHP-Pt-D3_48h | SRR34364288 | SAMN49784149 | 1.38E+08 | 20.77148 | 1.35E+08 | 19.04193 | 97.71603 | 93.36439 | 53.98158 | 67453114 | 58683366 | 87.0 |
